# Distinct representational structure and localization for visual encoding and recall during visual imagery

**DOI:** 10.1101/842120

**Authors:** Wilma A. Bainbridge, Elizabeth H. Hall, Chris I. Baker

## Abstract

During memory recall and visual imagery, reinstatement is thought to occur as an echoing of the neural patterns during encoding. However, the precise information in these recall traces is relatively unknown, with previous work primarily investigating either broad distinctions or specific images, rarely bridging these levels of information. Using ultra-high-field (7T) fMRI with an item-based visual recall task, we conducted an in-depth comparison of encoding and recall along a spectrum of granularity, from coarse (scenes, objects) to mid (e.g., natural, manmade scenes) to fine (e.g., living room, cupcake) levels. In the scanner, participants viewed a trial-unique item, and after a distractor task, visually imagined the initial item. During encoding, we observed decodable information at all levels of granularity in category-selective visual cortex. In contrast, information during recall was primarily at the coarse level with fine level information in some areas; there was no evidence of mid-level information. A closer look revealed segregation between voxels showing the strongest effects during encoding and those during recall, and peaks of encoding-recall similarity extended anterior to category-selective cortex. Collectively, these results suggest visual recall is not merely a reactivation of encoding patterns, displaying a different representational structure and localization from encoding, despite some overlap.

## Introduction

When we visually recall an object or scene, our memory contains rich object and spatial information (Bainbridge et al. 2019). During such recollection, our brain is thought to reinstate neural patterns elicited by the initial perception (McClelland et al. 1995; Buckner and Wheeler 2001; Tompary et al. 2016; Dijkstra et al 2019). One common view is that the hippocampus indexes populations of neocortical neurons associated with that memory (Teyler and Rudy 2007; Danker and Anderson 2010; Schultz et al. 2019). Under this view, representations in hippocampus are largely independent of a memory’s perceptual content (Davachi 2006; Liang et al. 2013; Huffman and Stark 2014). In contrast, the neocortex is thought to show sensory reinstatement, where the same regions show the same representations during recall as during encoding (Wheeler et al. 2000; Kahn et al. 2004; Staresina et al. 2012; Ritchey et al. 2013; Lee et al. 2012; O’Craven and Kanwisher 2000; Dijkstra et al. 2017). However, prior work has focused on specific levels of information (e.g. broad stimulus class, specific image) and the extent to which representations during recall reflect the same information as during perception, at all levels of granularity (from individual exemplar up to broad stimulus category), is unclear. Here, using ultra-high-field (7T) fMRI, we conducted an in-depth investigation of the content of encoded and recalled representations of objects and scenes across cortex, hippocampus, and the medial temporal lobe, assessing the granularity of detail in the representations of individual items.

First, we employed a hierarchically organized stimulus set (Figure 1a) with three levels of granularity from coarse (scenes/objects) to mid (e.g., natural/manmade scenes) to fine (e.g., bedrooms/conference rooms) level. Prior work comparing encoding and recall have primarily investigated memory content at opposite ends of this granularity spectrum. At a coarse level, recall of stimulus classes (faces, scenes, objects) have been reported to reactivate high-level visual regions (Polyn et al. 2005; Johnson et al. 2009; Reddy et al. 2010; LaRocque et al. 2013) and produce differentiable responses in hippocampus (Ross et al 2018). At the fine level, other work has shown reinstatement for individual images, with specific visual stimuli decodable in high-level visual cortex (Dickerson et al. 2007; Buchsbaum et al. 2012; Lee et al. 2012; Kuhl and Chun 2014) and medial temporal lobe (Zeineh et al. 2003; Gelbard-Sagiv et al. 2008; Chadwick et al. 2010; Wing et al. 2015; Mack and Preston 2016; Tompary et al. 2016; Lee et al. 2019).

**Figure 1.**
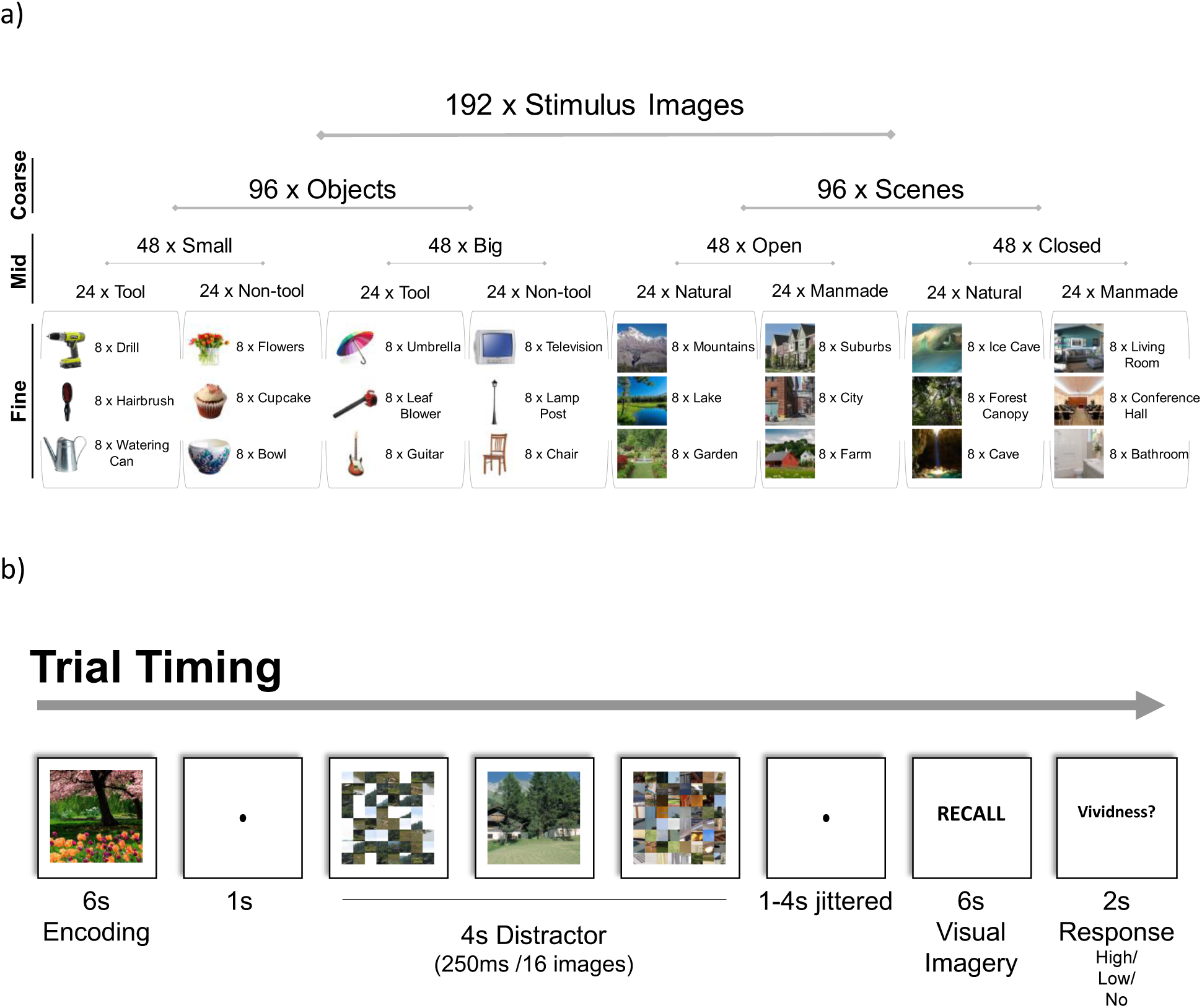
Experimental stimuli and task. (a) Nested structure of stimuli and example images. 192 trial-unique images were encoded and recalled by participants, arranged under nested structure based on the *coarse* level (object / scene), *mid* level (e.g., open / closed or natural / manmade scene), and *fine* level (mountains / lake) of stimulus organization. Each fine-level category contained 8 different exemplar images (e.g., 8 different lake photographs). (b) The timing of each trial. Participants studied an image for 6 s, performed a distractor task requiring detection of an intact image amongst scrambled images for 4 s, and then after a randomized jitter of 1-4 s, recalled the original image through visual imagery for 6 s. Finally, they indicated the vividness of their memory with a button press.

Decoding for specific images (Thirion et al. 2006; Naselaris et al. 2015), positions (Stokes et al. 2011) and orientations (Klein et al. 2004; Albers et al. 2013) is even present in early visual cortex during visual imagery. However, it is often unclear what information is driving discrimination across the brain: fine-level image-specific information, coarse-level perceptual category information, or information unrelated to stimulus content such as memory strength. For example, while recalled grating orientation is decodable from early visual cortex (V1-V3), reinstatement strength but not content is decodable from the hippocampus (Bosch et al. 2014). Further, few studies have investigated the ability to detect reinstatement of mid-level information (e.g., is it a natural or manmade scene, a big or small object) during recall, even though such information is known to be decodable during perception (e.g., Park et al. 2011; Kravitz et al. 2011; Konkle et al. 2012). Our approach using nested levels of stimulus information reveals what granularity of information is contained in regions across the visual processing pathway, and whether reinstatement is simply an echo of the same response from encoding to recall.

Second, to isolate the activity specific to recall, we adopted a visual imagery task focusing on recall of individual items without requiring the learning of cue-stimulus associations, which have commonly been used (e.g., Ganis et al. 2004; Kuhl et al. 2012; Zeidman et al. 2015a; Jonker et al. 2018). Recalled representations in associative tasks are likely to contain information not only about the recalled item, but also the cue and the association itself. Further, there are differences in neocortex when performing an associative versus item-based memory task (Staresina and Davachi 2006). In fact, the neural representation of a target may be largely dependent on what cue it is associated with (Xiao et al. 2017). Here, we employ an item-based recall task in which participants encode trial-unique images, and following a distractor task, recall that specific image. This approach allows us to investigate the recall of individual items, without the learning of associations.

Using this direct recall task and nested stimulus structure, we find striking differences in the representational structure and spatial localization for visual encoding and recall, suggesting recall patterns are not just a repetition of patterns during encoding, despite some similarities.

## Materials and Methods

### Participants

Thirty-four adults were recruited for the experiment. All participants were healthy, right-handed, and had corrected or normal vision. Twelve participants were unable to complete the experiment due to discomfort in the 7T scanner, drowsiness, or scanner malfunction, and their data were excluded from the study. This level of participant dropout is not unusual for 7T scans, given that nausea and vertigo occasionally occur, and the bore is more restrictive than the more standard 3T scanners. The final set of participants included twenty-two adults (fifteen female; mean age: 24 years, standard deviation: 3.4 years, range: 19-35 years). All participants provided consent following the guidelines of the National Institutes of Health (NIH) Institutional Review Board (National Institute of Mental Health Clinical Study Protocol NCT00001360, 93M-0170), and were compensated for their participation.

### Stimuli

Stimulus images comprised 192 images with nested categorical structure (Figure 1a). We refer to the different levels of information as coarse, mid, and fine. At a coarse level, 50% of the stimuli were objects and 50% scenes.

At a mid-level, the objects and scenes were varied according to factors known to show differential responses in the brain during perception. The objects were made up of four object types, varying along two factors: 1) small / big objects (Konkle et al. 2012), and 2) tool / non-tool objects (Valyear et al. 2007; Mahon et al. 2007; Beauchamp and Martin 2007). Big objects were selected as objects generally larger than a 1-foot diameter and small objects were those smaller than a 1-foot diameter. Tools were defined as objects commonly grasped by one’s hands using a power grip (e.g., Grèzes et al. 2003), although note that there are multiple ways tools are defined in the field (Lewis 2006). Similarly, the scenes were made up of four scene types, varying along two factors: 1) natural / manmade (Park et al. 2011), and 2) open / closed (Kravitz et al. 2011). Natural scenes were defined as those primarily made up of natural objects (i.e., plants, rocks, sand, ice), while manmade scenes were primarily made of artificial objects (i.e., buildings, furniture). Open scenes were defined as those with an open spatial extent, while closed scenes were defined as those in which the viewer is enclosed by boundaries (Park et al. 2011).

At a fine-level, each object or scene type contained three categories, with eight exemplars for each object or scene category (e.g., <u>small, non-tool objects</u>: bowl, cupcake, flowers; <u>closed, manmade scenes</u>: bathroom, conference hall, living room; see Figure 1a for all fine-level categories). Images were all square 512 x 512 pixel images presented at 8 degrees of visual angle, and objects were presented cropped in isolation on a white background.

### In-scanner recall task and post-scan recognition task

Participants first completed a single run of a 7-min 6-sec block-design localizer scan to identify scene- and object-selective regions. In this localizer, participants viewed 16-sec blocks of images of objects, scenes, faces, and mosaic-scrambled scenes and identified consecutive repeated images. All images used in the localizer were distinct from those used in the main experiment. Participants then completed eight runs of an item-based memory recall task requiring visual imagery (Figure 1b). In each trial, participants studied a trial-unique stimulus image for 6 s. After a 1 s fixation, they performed a distractor task in which they viewed a stream of 16 quickly presented images (250 ms each) and had to press a button as soon as they saw the sole intact image in a stream of mosaic-scrambled images. Scrambled and intact target images were taken from a separate stimulus set, and were chosen to be of the same coarse level (i.e., object or scene) as the studied image in order to keep general visual properties consistent (i.e., not switching from one type of stimulus to another). These distractor images were taken with random mid and fine levels, unrelated to the stimulus being encoded and recalled. Intact object images were presented as an intact object against a mosaic-scrambled background, so that participants would have to fixate the object to successfully perform the task (rather than identify white edges). The distractor task lasted for 4 s total and was followed by a 1-4 s jittered interval in which participants were instructed to wait and maintain fixation. The word “RECALL” then appeared on the screen for 6 s, and participants were instructed to silently visually imagine the originally studied image in as much detail as possible. Finally, following the “RECALL” phase, participants were given 2 s to press a button indicating the vividness of their memory as either no memory, low vividness, or high vividness. The next trial then continued after a 1 s delay. Participants were instructed that the task was difficult, and they should focus on reporting their vividness truthfully. On average, participants reported ‘high vividness’ on 60.8% of trials (SD=16.9%), ‘low vividness’ on 29.9% of trials (SD=12.8%), and ‘no memory’ on 9.31% of trials (SD=8.38%). Trials in which participants indicated “no memory” were not included in any of the main analyses of the data. Each run contained 24 trials, lasting 8 min 38 s, and participants completed 8 runs total. Each run included three “catch trials” that skipped the recall phase, in order to keep participants vigilant, to discourage them from pre-emptively recalling the target image, and to better separate encoding from distractor and recall phases during deconvolution. Each fine-level stimulus category (e.g., guitar, cupcake) was shown once per run, and each stimulus exemplar image was only used once in the entire experiment, so that there would be no memory effects on subsequent presentations of the same image.

After the scan, participants performed a post-scan recognition task to test their memory for the images studied in the scanner. Participants were presented with all 192 images studied in the scanner randomly intermixed with 192 foil images of the same fine-level stimulus categories and were asked to indicate for each image whether it was old or new. Two participants were unable to complete the post-scan recognition task due to time constraints. Analyses on the post-scan recognition data as well as vividness ratings are reported in the Supplementary Material (SM1, SM2).

### MRI acquisition and preprocessing

The experiment was conducted at the NIH, using a 7T Siemens MRI scanner and 32-channel head coil. Whole-brain anatomical scans were acquired using the MP2RAGE sequence, with 0.7 mm isotropic voxels. Whole-brain functional scans were acquired with a multiband EPI scan of in-plane resolution 1.2 x 1.2 mm and 81 slices of 1.2 mm thickness (multiband factor = 3, repetition time = 2 s, echo time = 27 ms, matrix size = 160 × 160, field of view = 1728 × 1728, flip angle = 55 degrees). Slices were aligned parallel with the hippocampus and generally covered the whole brain (when they did not, sensorimotor parietal cortices were not included). Functional scans were preprocessed with slice timing correction and motion correction using AFNI and surface-based analyses were performed using SUMA (Cox 1996; Saad and Reynolds 2012).

### FMRI Region of Interest (ROI) Definitions

For each participant, key ROIs for early visual cortex, object selective cortex, scene selective cortex, and hippocampus were determined *a priori* and defined using functional and anatomical criteria (Figure 2). Using the independent functional localizer, we identified three scene-selective regions with a univariate contrast of scenes > objects: PPA (Epstein & Kanwisher 1998), medial place area (MPA; Silson et al. 2016), and occipital place area (OPA; Dilks et al. 2013). We localized object-selective regions lateral occipital (LO) and posterior fusiform (pFs) with a univariate contrast of objects > scrambled images (Grill-Spector et al. 2001). Finally, we localized early visual cortex (EVC) with a univariate contrast of scrambled images > baseline. With each contrast, the functional ROIs were defined as the contiguous set of at least 20 voxels showing significant activation for the contrast, located within the broader anatomical areas described in the literature (e.g., PPA should be in and around the collateral sulcus, Epstein and Baker 2019; LO should be within lateral occipital regions). For all contrasts, we first identified these contiguous sets of voxels with a univariate contrast with a False Discovery Rate (FDR)-corrected threshold of *q*=0.001. When a contiguous set of voxels could not be identified, we looked at increasingly liberal thresholds of *q*=0.005, *q*=0.01, *q*=0.05, *p*=0.001, *p*=0.005, *p*=0.01, and *p*=0.05 until a contiguous set of 20 voxels passing that threshold was identified. If no contiguous set of voxels was identified at this threshold, then the ROI was determined missing for that given participant. Left and right ROIs were combined to create bilateral ROIs in the analyses. Overlapping voxels between scene- and object-selective regions were discarded from any ROI. LO, pFs, PPA, and EVC were identified in 22 participants, OPA in 21 participants, and MPA in 20 participants. Anatomical ROIs were localized using FreeSurfer’s recon-all function using the hippocampal-subfields-T1 flag (Iglesias et al. 2015), and then visually inspected for accuracy. This hippocampus parcellation function splits the hippocampus into the head/body (Hip-HB) and tail (Hip-T), and within the head/body region further segments the hippocampus into different subfields (dentate gyrus, CA1, CA3, and subiculum; Iglesias et al. 2015). We did not find meaningful differences across subfields (all subfields either showed identical results to the Hip-HB or Hip-T), but report those results in the Supplementary Material (SM3). This FreeSurfer parcellation also localized the perirhinal cortex (PRC) and parahippocampal cortex (PHC) within the medial temporal lobe (MTL). PHC was determined as a participant’s anatomically defined PHC minus voxels already contained with their functionally defined PPA. For the main body of the text, we report the results from the Hip-HB (with subfields combined), Hip-T, PRC, and PHC. A table of ROI sizes by participant is provided in the Supplementary Material (SM4).

**Figure 2.**
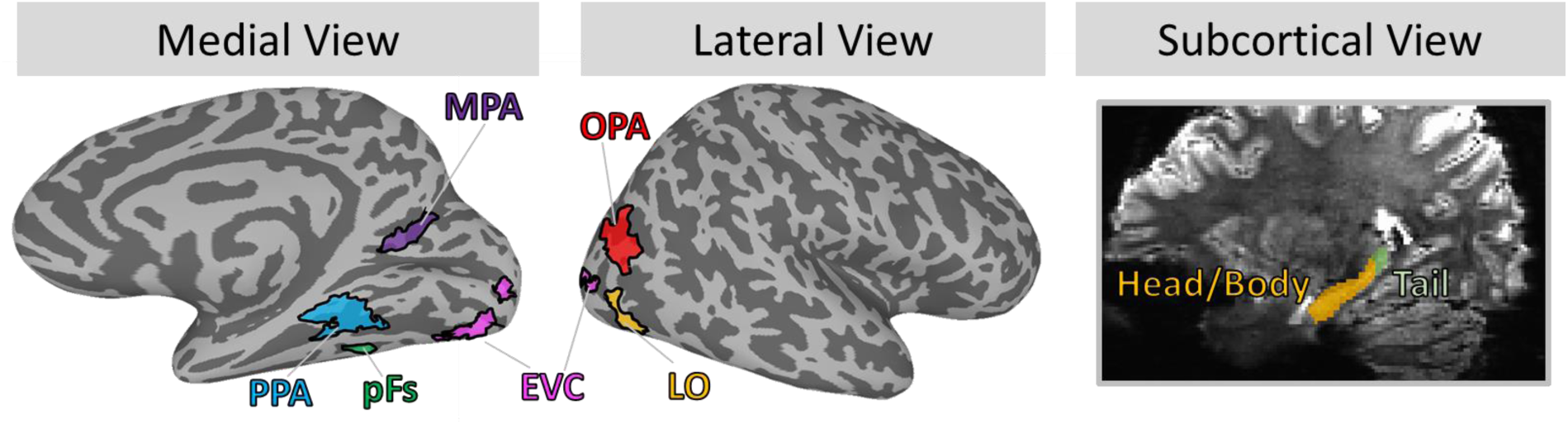
Main regions of interest (ROI). The current study focused on a set of visual and memory-related ROIs. Visual regions consisted of early visual cortex (EVC), object-selective regions of the lateral occipital (LO) and the posterior fusiform (pFs), and scene-selective regions of the parahippocampal place area (PPA), medial place area (MPA), and occipital place area (OPA). Visual regions were individually localized using functional localizers in each participant; shown here are probabilistic ROIs of voxels shared by at least 12% of participants. Memory-related regions consisted of the hippocampus divided into anterior (head and body) and posterior (tail) subregions, as well as the perirhinal cortex (PRC, not shown) and parahippocampal cortex (PHC, not shown). These ROIs were segmented automatically using anatomical landmarks.

### Whole-Brain Univariate Analyses

We conducted whole-brain univariate contrasts using a general linear model (GLM) that split the trials into six regressors along two factors: 1) encoding / distractor / recall, and 2) scenes / objects. Six additional regressors for movement were also included. Additionally, trials in which participants indicated they had “no memory” for the item were modeled separately as three regressors (for the encoding, distractor, and recall periods) in the GLM to avoid them contributing to either target stimulus responses or an implicit baseline. We then performed whole-brain *t*-contrasts of scenes vs. objects separately during encoding and recall. All whole-brain contrasts were projected onto the cortical surface using AFNI surface mapper SUMA (Saad and Reynolds 2012).

With these whole-brain analyses, we located and compared the peak voxels of activation during encoding and recall. For each participant, we localized the peak voxel within each broad visual ROI definition (e.g., for PPA, voxels in and around the collateral sulcus, Epstein and Baker 2019) separately for encoding and recall. We extracted the MNI coordinates for each participant for the voxels with the highest object activation near LO and pFs, and the highest scene activation near PPA, MPA, and OPA. The peaks of encoding and recall were directly compared across participants with a paired Wilcoxon signed rank test, comparing the median anterior-posterior coordinates between encoding and recall.

### Representational Similarity Analyses and Discrimination Indices

Multivariate analyses were conducted to look at the representations of different stimulus information during encoding and recall, across brain regions. For these analyses, the experimental data were first split into two independent halves – even runs and odd runs. For each split half, a GLM was calculated, modeling separate regressors for each fine-level category (e.g., cupcake) for the encoding period (6 s boxcar function), distractor period (4 s boxcar), and recall period (6 s boxcar). Each event (e.g., encoding a cupcake) thus had two resulting beta estimates: one across even runs, and one across odd runs. As in the univariate analysis GLM, trials in which participants indicated they had no memory for the image were captured with three additional regressors for the encoding period, the distractor period, and the recall period. The estimated motion parameters from the motion correction were included as six further regressors.

We then investigated the similarity between different types of stimulus information during encoding, recall, and distractor periods, using representational similarity analyses (RSA; Kriegeskorte et al. 2008). For each ROI, we created a representational similarity matrix (RSM) comparing the similarity of all pairs of fine-level stimulus category (e.g., cupcake vs. guitar). Similarity was calculated as the Pearson’s correlation between the voxel values (t-statistic) in an ROI for one fine-level category (e.g., cupcake) from one half of the runs (e.g., odd runs), with the voxel values for another category (e.g., guitar) from the other half (e.g., even runs). Specifically, pairwise item similarity was taken as the average of the correlation with one split (odd runs for item A, even runs for item B) and correlation with the opposite split (even runs for item A, odd runs for item B). This metric indicates the similarity in the neural representations of two categories, and importantly, because the comparisons use separate halves of the data, we can observe a category’s similarity to itself across runs. This self-similarity measure thus quantifies the degree to which a given region shows similarity across exemplars within category (i.e., are cupcakes similar to other cupcakes). Correlation coefficients were all corrected with Fisher’s Z-transformations. We focused our main analyses on three RSMs: 1) correlations of the encoding responses (Encoding RSM), 2) correlations of the recall responses (Recall RSM), 3) correlations of the encoding responses with the recall responses (Cross-Discrimination RSM). These different classifications allow us to see what stimulus information exists separately during encoding and recall, as well as what information is shared between encoding and recall.

From these RSMs, we conducted discriminability analyses, which show the degree to which each ROI can discriminate the different conditions of fine-, mid-, and coarse-level information (e.g., do the responses in PPA discriminate natural vs. manmade scenes?). For each comparison of interest, we computed a discrimination index *D,* calculated as the difference of the mean across-condition correlations (e.g., scenes with objects) from mean within-condition correlations (e.g., scenes with other scenes; Kravitz et al. 2011; Cichy et al. 2014; Harel et al. 2013; Harel et al. 2014; Henriksson et al. 2015). The intuition behind this index is that if an ROI contains information about that comparison, then within-condition similarity should be higher than across-condition similarity (e.g., if the PPA *does* discriminate natural vs. manmade scenes, then natural scenes should be more similar to other natural scenes than manmade scenes). Discriminability analyses at all levels of stimulus granularity were calculated from the same underlying correlation matrix, and there were close to the same number of trials contributing to the calculation of each cell in the matrix (only differing due to the exclusion of no-memory trials). However, do note that the comparisons of different granularity use different proportions of the matrix; e.g., the coarse level of objects versus scenes utilizes the whole matrix, while the mid level of natural versus manmade only looks within scenes. We compared these discrimination indices versus a null hypothesis of 0 discrimination using one-tailed t-tests. While multivariate analyses may often violate the assumptions of parametric statistics (Allefeld et al. 2016), in practice, one-tailed t-tests to evaluate discrimination indices are not meaningfully different from non-parametric methods (Nili et al. 2020). However, we confirmed all results hold when also calculating significance with a permutation test across 1,000 RSM permutations (Supplementary Material SM5). Mid-level discriminability was only computed within same-coarse-level items (e.g., only scenes were used for the natural vs. manmade comparison), and fine level discriminability was only computed within same-mid-level items (e.g., when looking at the discriminability of living rooms, they were only compared to other closed, manmade scenes). Refer to Figure 3 for a depiction of these discrimination indices and to see example RSMs. All statistics reported are FDR-corrected within each ROI across all 21 discriminations (the seven discriminations shown in Figure 3, each for encoding, recall, and cross-discrimination) at a value of *q* < 0.05.

**Figure 3.**
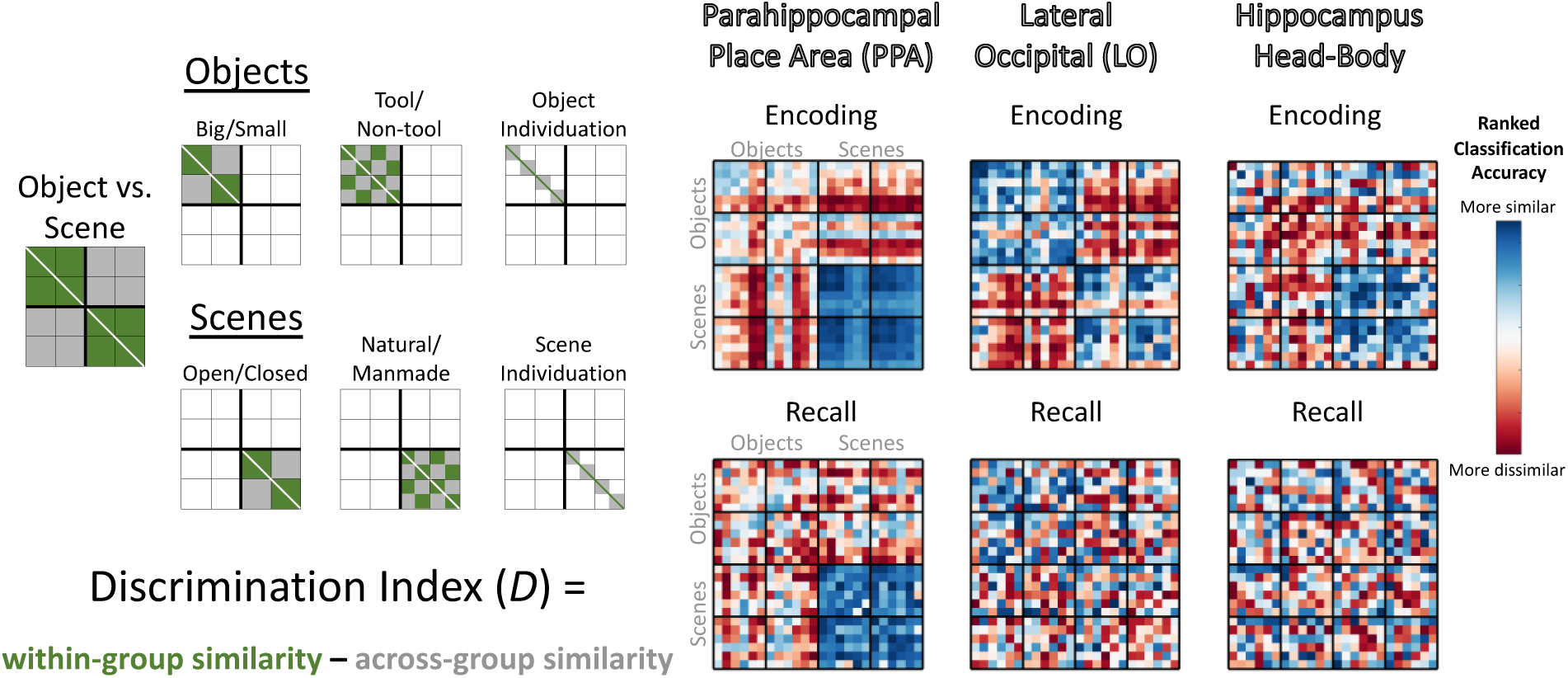
Calculating information discriminability from representational similarity matrices. (Left) Depictions of the cells of the representational similarity matrices (RSMs) used to calculate discrimination indices for key regions of interest (ROIs). The RSMs represent pairwise Pearson’s correlations of stimulus groupings calculated from ROI voxel t-values, compared across separate run split halves (odd versus even runs). These depictions show which cells in the matrices are used in the calculation of discriminability of different properties, with green cells indicating within-condition comparisons, which are compared with grey cells indicating across-condition comparisons. For all discriminability calculations except fine-level discrimination of individual categories, the diagonal was not included. All operations were conducted on the lower triangle of the matrix, although both sides of the diagonal are shown here for clarity. (Right) Examples of encoding and recall RSMs from the data in the current study, specifically the rank-transformed average RSM for the parahippocampal place area (PPA), lateral occipital (LO), and the hippocampus head and body. Blue cells are more similar, while red cells are more dissimilar.

### Discrimination-based Searchlight Analyses

We also conducted discriminability analyses using spherical searchlights (3-voxel radius) in two ways. First, we conducted discriminability analyses (as described above) for searchlights centered on voxels in the ROIs. For each searchlight, we obtained a scene-object discriminability metric during encoding and one during recall, allowing us to examine the relationship between encoding and recall information in these ROIs. Note that while the center voxel in the searchlight was located within each given ROI, peripheral voxels could fall outside of an ROI’s boundaries. This is to ensure that searchlights are of equal volume throughout the ROI and will result in only a small amount of smoothing of the ROI’s borders (e.g., 3 voxels at maximum).

Second, we conducted discriminability analyses in searchlights iteratively moved through each individual’s brain, to examine ability to discriminate information outside of our pre-defined ROIs. Group maps were combined with a one-tailed t-test comparing group discrimination indices versus no discrimination (0). Group maps were thresholded at *p* < 0.005 uncorrected for visualization purposes however we also provide unthresholded maps. We conducted these searchlights looking at both discriminability of information within memory process type (encoding or recall), as well as ability to cross-discriminate information between encoding and recall. We also identified the locations of peak voxels for encoding, recall, and cross-discrimination, using the same methods as described in the *Whole-Brain Univariate Analyses*.

### Encoding-recall correlation and overlap analyses

In order to directly compare encoding and recall information within ROIs, we conducted two separate analyses. We specifically focused on coarse level discrimination of objects versus scenes, as this discrimination is reliably found across regions for both encoding and recall (see Results).

First, we calculated the correlation between encoding and recall discrimination indices in the searchlights within each ROI (see previous section). For each searchlight centered within an ROI, we Spearman rank correlated its coarse level discrimination index (scenes vs. objects) between encoding and recall. This analysis reveals the degree to which voxels that represent encoding information also represent recall information. High correlations indicate that voxels that can discriminate objects versus scenes during encoding can also discriminate them during recall, while low correlations provide evidence for no relationship between encoding and recall discriminability. Significance was calculated using a non-parametric Wilcoxon signed rank test, comparing the rank correlations against a null median of 0.

Second, given the relatively low correlations we observed, we conducted an overlap analysis to determine the degree to which the most discriminative voxels are the same between encoding and recall. To perform this analysis, for each ROI, we took the top 10% discriminating encoding voxels and compared their overlap with the top 10% discriminating recall voxels. Chance level of overlap was calculated with permutation testing, by taking two random sets of searchlights (rather than the top ranked searchlights) consisting of 10% of the ROI size. Across 100 permutations per ROI per participant, we calculated the overlap between these two shuffled sets, and then took the average across all permutations as the chance level for each participant. This permuted level of chance ultimately resolves to 10% across all ROIs, which matches the computed chance level for this analysis - if you take two random sets of 10% of voxels, by chance, 10% of those voxels should overlap. Significance was calculated with a non-parametric paired Wilcoxon rank sum test comparing the true overlap percentage with the permuted random overlap percentage.

## Results

In the following sections, we examine the relationship between representations elicited during encoding and recall. First, we compare granularity of stimulus content representations in object- and scene-selective visual ROIs and the hippocampus. We observe reduced information during recall, particularly for mid-level information. Second, to directly compare encoding and recall representations, we conduct searchlight analyses to investigate the distribution of voxels showing the strongest discrimination during encoding, recall and cross-discriminability between these two phases, both within and outside the ROIs. We observe little correlation between discrimination during encoding and recall and find that the voxels that represent recalled information are frequently distinct from those that represent encoding information, with the strongest representations during recall anterior to the category-selective regions traditionally studied during perception.

### Decoding stimulus content from scene- and object-selective visual regions and medial temporal lobe

What aspects of a visual memory are represented in scene- and object-selective areas and medial temporal lobe during encoding and recall? We asked this question by discriminating stimulus information from the patterns of blood oxygen level dependent (BOLD) responses at various scales of stimulus granularity, ranging from a coarse level (scenes, objects), to a mid-level (e.g., natural/manmade scene, big/small object), to a fine level (e.g., cupcake, guitar). This discrimination was conducted across independent exemplars, never including the same images in the training and testing sets of the decoding model. This allowed us to see what levels of information are represented in these regions, separate from an ability to distinguish identical images. Discrimination indices and their corresponding *p*-values (see Methods) for all ROIs are reported in Supplementary Material SM6. Here, in the text we only describe statistics that pass FDR correction, but all values including those where *p* < 0.05 but *p* does not pass FDR correction are included in this table.

### Visual ROIs: Detailed information during encoding, limited information during recall

We investigated discriminability in object-selective regions LO and pFs, scene-selective regions PPA, MPA, and OPA, and early visual cortex (Figure 4, refer to Supplementary Material SM6 for discrimination indices and individual statistics).

**Figure 4.**
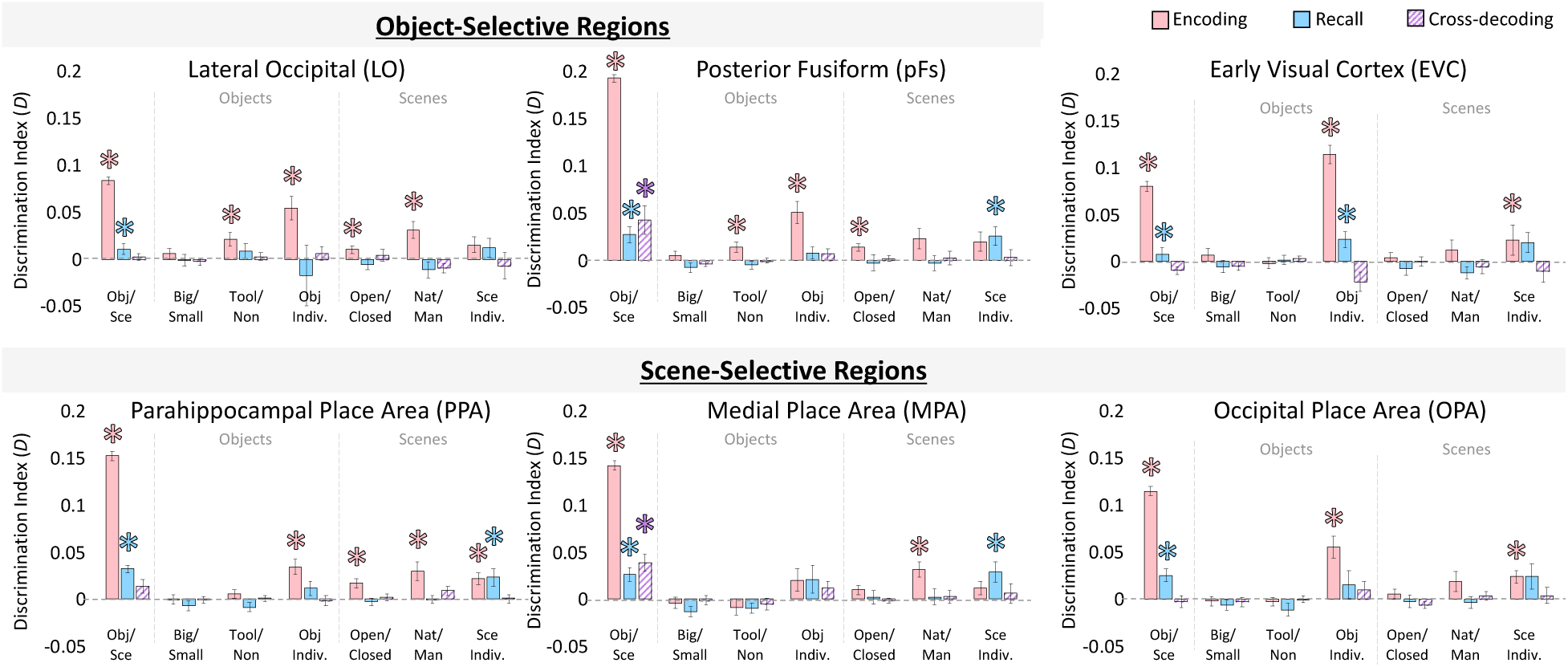
Information discriminability in scene- and object-selective regions. Discriminability for visual regions of interest (ROIs) for each stimulus property was calculated from the RSMs (as in Figure 3). Bar graphs indicate mean discrimination index for different comparisons across ROIs, are split by coarse stimulus class, and show three levels of discrimination: 1) the coarse level (objects versus scenes), 2) the mid level (objects: big/small, tools/non-tools; scenes: open/closed, natural/manmade), and 3) the fine level (specific object and scene categories). The y-axis represents the average discrimination index (*D*), which ranges from -1 to 1. Significance (*) indicates results from a one-tailed t-test versus 0, with a FDR-corrected level of *q* < 0.05 (applied to all 21 comparisons within each ROI). Values that do not pass FDR correction can still be seen in Supplementary Material SM6. Pink bars indicate discriminability during encoding trials, blue bars indicate discriminability during recall trials, and hatched purple bars indicate cross-discriminability (i.e., there is a shared representation between encoding and recall). Error bars indicate standard error of the mean.

We first examined what information was discriminable during the encoding period. All object- and scene-selective regions could discriminate coarse level information (objects vs. scenes), all *p* < 10^-4^. For mid-level object information, tool/non-tool could be discriminated in object-selective regions LO (*p =* 0.009) and pFs (*p* = 0.012), but object size did not show significant discriminability in any region. For mid-level scene information, open/closed could be discriminated in LO (*p* = 0.009), pFs (*p* = 0.002), and PPA (*p* = 0.001), while manmade/natural could be discriminated in LO (*p* = 0.001), PPA (*p* = 0.003), and MPA (*p* = 4.76 × 10^-4^). Finally, fine-level object information could be discriminated in all regions except MPA (all *p* < 0.001), while the fine level for scenes could be discriminated in scene-selective regions PPA (*p* = 7.64 × 10^-4^) and OPA (*p* = 0.002). In addition, response patterns in the encoding period for all visual ROIs was predictive of reported memory vividness, and patterns in the LO, PPA, and OPA were predictive of subsequent recognition (Supplementary Material SM1, SM2). Overall, these results confirmed the findings of prior studies (e.g., Valyear et al. 2007; Walther et al. 2009; Park et al. 2011; Kravitz et al. 2011; Troiani et al. 2012), in which during encoding and perception, responses in scene- and object-selective regions can be used to distinguish various levels of information about visually presented scenes and objects.

We next investigated the information present in these ROIs during recall. Discrimination of coarse-level information was significant in all visual regions (all *p* < 0.001). However, no region showed significant discriminability for any mid-level information (big/small, tool/non-tool for objects and open/closed, natural/manmade for scenes; all *p* > 0.10). Also, no region showed fine-level object information during recall. However, significant fine-level information during recall of scenes was present in pFs (*p* = 0.009) as well as scene regions PPA (*p* = 0.011) and MPA (*p* = 0.008). In addition, response patterns from all visual areas during recall were predictive of recall vividness, although not predictive of subsequent recognition (Supplementary Material SM1, SM2). These results reveal that while visual regions maintain coarse-level information during recall, we find no evidence for mid-level stimulus information. Despite the lack of mid-level information, however, there is fine-level information in some regions.

To investigate which regions show a shared neural representation during encoding and recall, we conducted a cross-discrimination analysis identifying the degree to which a region shows similar patterns between encoding and recall. The only significant cross-discrimination was for the coarse level (objects versus scenes), which was found in pFs (*p* = 0.006) and MPA (*p* = 1.59 × 10^-4^). Significant cross-discrimination did not emerge for any mid-level information in any ROI (all *q* > 0.05), nor at the fine level in any ROI (all *q* > 0.05). These findings imply that encoding and recall may differ in their representational structure across different levels of information.

Given these effects in object- and scene-selective regions, we conducted a follow-up analysis to look at visual responses outside of category selective cortex, namely early visual cortex (EVC; Figure 4, Supplementary Material SM6). During encoding, EVC showed significant discrimination at the coarse level (*p* = 4.43 × 10^-7^), but no discrimination at the mid-level for scenes or objects (all *q* > 0.05). However, EVC did show significant discrimination of the fine-level for both objects (*p* = 5.13 × 10^-7^) and scenes (*p* = 0.005). During recall, EVC again showed significant coarse-level discrimination (*p* = 0.004), no significant mid-level discrimination (all *p* > 0.10), and significant fine-level discrimination for objects (*p* = 0.008) although not for scenes (*p* > 0.05). EVC did not show significant cross discrimination at any level (all *p* > 0.20). These results suggest that retinotopic information—driven by the visual features of different object categories and their differences from scenes—is likely discriminable during recall. However, mid-level information did not show differences in early visual processing during encoding, and was not discriminable during recall.

### Hippocampus and Medial Temporal Lobe Show Coarse Level Information During Encoding

We conducted the same analyses in the hippocampus and medial temporal lobe regions (MTL) consisting of the perirhinal cortex (PRC) and parahippocampal cortex (PHC) (Figure 5). We primarily focused on the segregation of the hippocampus into anterior (Hip-HB) and posterior (Hip-T) regions, but results for the individual subfields can be found in the Supplementary Material (SM3).

**Figure 5.**
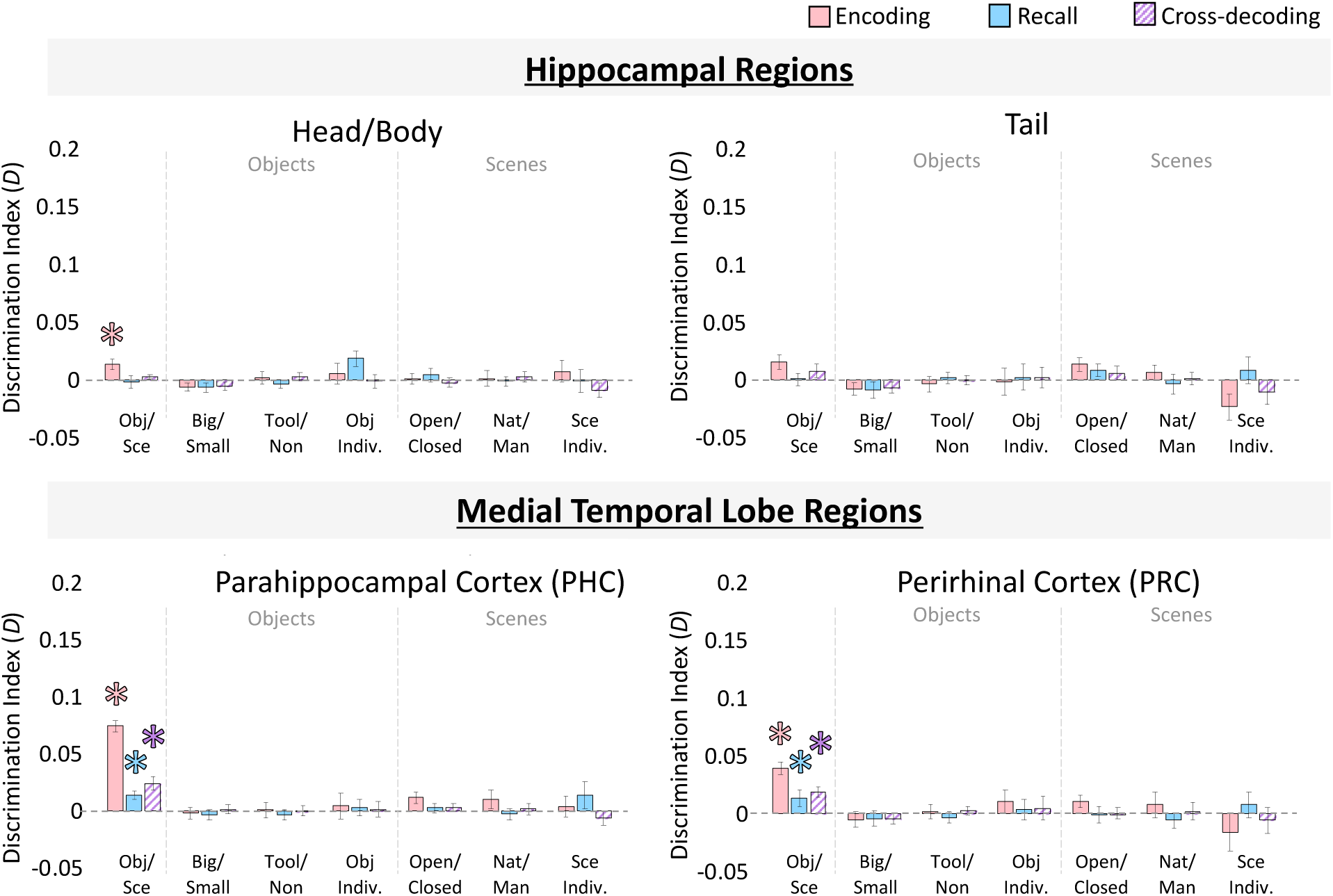
Information discriminability in the hippocampus and medial temporal lobe. Discriminability for hippocampal ROIs, perirhinal cortex (PRC), and parahippocampal cortex (PHC) for each stimulus property was calculated from the RSMs. Bar graphs are displayed in the same manner as Figure 4, and indicate mean discrimination index for comparisons of different levels of stimulus information (coarse, mid-, and fine levels for objects and scenes). Pink bars indicate discriminability during encoding trials, blue bars indicate discriminability during recall trials, and hatched purple bars indicate cross-discriminability (i.e., there is a shared representation between encoding and recall). Error bars indicate standard error of the mean. Asterisks (*) indicate significance at a FDR corrected level of *q* < 0.05.

During encoding, significant coarse-level discrimination of objects versus scenes was present in Hip-HB (*p =* 3.28 × 10^-4^), PRC (*p =* 5.74 × 10^-5^), and PHC (*p =* 4.62 × 10^-6^), but not Hip-T (*p* = 0.108). There was no mid-level information present in any of these regions (all *q* > 0.05), nor was there fine-level information (all *q* > 0.05). During recall, course-level information was not detected in the hippocampus (Hip-HB: *p* = 0.82; Hip-T: *p* = 0.48), but was discriminable in PRC (*p* = 0.004) and PHC (*p* = 0.003). No mid- or fine-level information was present during recall in any of these regions (all *p* > 0.20). Finally, significant cross-discriminability between encoding and recall was found in the PRC (2.53 × 10^-4^) and PHC (3.99 × 10^-4^), but not in the hippocampus (Hip-HB: *p* = 0.082; Hip-T: *p* = 0.10). No mid-level or fine-level information was cross-discriminable in these regions (all *p* > 0.10).

Although hippocampus did not show content-related information beyond the coarse level during encoding, additional analyses revealed other discriminable information present in the hippocampus. Patterns during recall in the hippocampus were significantly predictive of reported memory vividness, although patterns during encoding were not (Supplementary Material SM1, SM2). Further, an analysis comparing the representational structure in different ROIs revealed that while visual areas were very dissimilar from the hippocampus during the encoding period, their patterns become more similar to those of the hippocampus during recall (Supplementary Material SM7, SM8).

### Discrimination of Information During the Distractor Period

To ensure any ability to discriminate information during recall was not due to bleed-over from the encoding period or active visual working memory strategies, we computed discrimination indices during the distractor period. Distractor stimuli differed by coarse level category (scenes, objects), and indeed coarse level information was available in LO (discrimination index *D* = 0.02, *p* = 5.94 × 10^-5^), PPA (*D* = 0.02, *p* = 5.00 × 10^-5^), and OPA (D = 0.02, *p* = 6.13 × 10^-5^), although not in pFs, MPA, Hip-HB, PRC, or PHC (*q* > 0.05), all of which showed discrimination during the encoding and recall periods (with the exception of the Hip-HB). Mid-level and fine level information was not discriminable in any ROI during the distractor period (*q* > 0.05), despite the presence of such information during the encoding period. Importantly, the lack of fine-level information during the distractor period contrasts with the stronger and significant fine-level information in pFs, PPA, and MPA during the recall period. Thus, it is highly unlikely that information measured during recall reflects carry over from the encoding period or active visual working memory strategies.

In sum, the analyses in this section reveal that while during encoding information can be discriminated in many of these ROIs from all levels of stimulus granularity (fine, mid, and course), there is limited information available during recall. Namely, while coarse- and fine-level information is available in many of the ROIs, mid-level information was not detected in any ROI. Significant cross-discrimination between encoding and recall was also only present at the coarse level of information. These results suggest distinct representational structure during encoding and recall, and motivate a direct comparison between encoding and recall representations at the sub-ROI level.

#### Direct comparison of encoding and recall discriminability within ROIs

We observed cross-discrimination in some regions (pFS, MPA, PRC, PHC), but not in others (LO, PPA, OPA, Hip-HB, Hip-T), providing mixed evidence for shared neural substrates for encoding and recall across these regions. To further investigate the relationship between encoding and recall discrimination, we directly compared discrimination indices across each region. Because only coarse-level information showed cross-discrimination in any region, we focused our analyses here on the coarse discrimination of objects versus scenes. For each ROI, we computed Spearman rank correlations between encoding and recall discrimination searchlights (see Methods). While some regions showed significant correlations between encoding and recall discriminability (MPA: Median Spearman’s rank correlation *ρ* = 0.210, Wilcoxon signed rank test: *Z* = 2.52, *p =* 0.012; LO: *ρ* = 0.104, *Z* = 3.17, *p =* 0.002; pFs: *ρ* = 0.200, *Z* = 2.26 *p =* 0.024; Hip-HB: *ρ* = 0.065, *Z* = 2.09, *p =* 0.036; PHC: *ρ* = 0.244, *Z* = 3.72, *p =* 2.01 × 10^-4^), others did not (PPA: *ρ* = -0.006, *Z* = 1.31, *p* = 0.189; OPA: *ρ* = 0.030, *Z* = 1.28, *p =* 0.200; Hip-T: *ρ* = -0.045, *Z* = 0.02, *p =* 0.987; PRC: *ρ* = 0.168, *Z* = 1.79, *p =* 0.074). However, even in those cases where we found significant effects, the correlations tended to be weak, and every ROI had 22% (5 out of 22) or more of the participants who showed negative correlations between encoding and recall. Moreover, the distribution of the plotted data often revealed an L-shape distribution with greatest similarity between encoding and recall for the voxels with the lowest discrimination scores (Figure 6).

**Figure 6.**
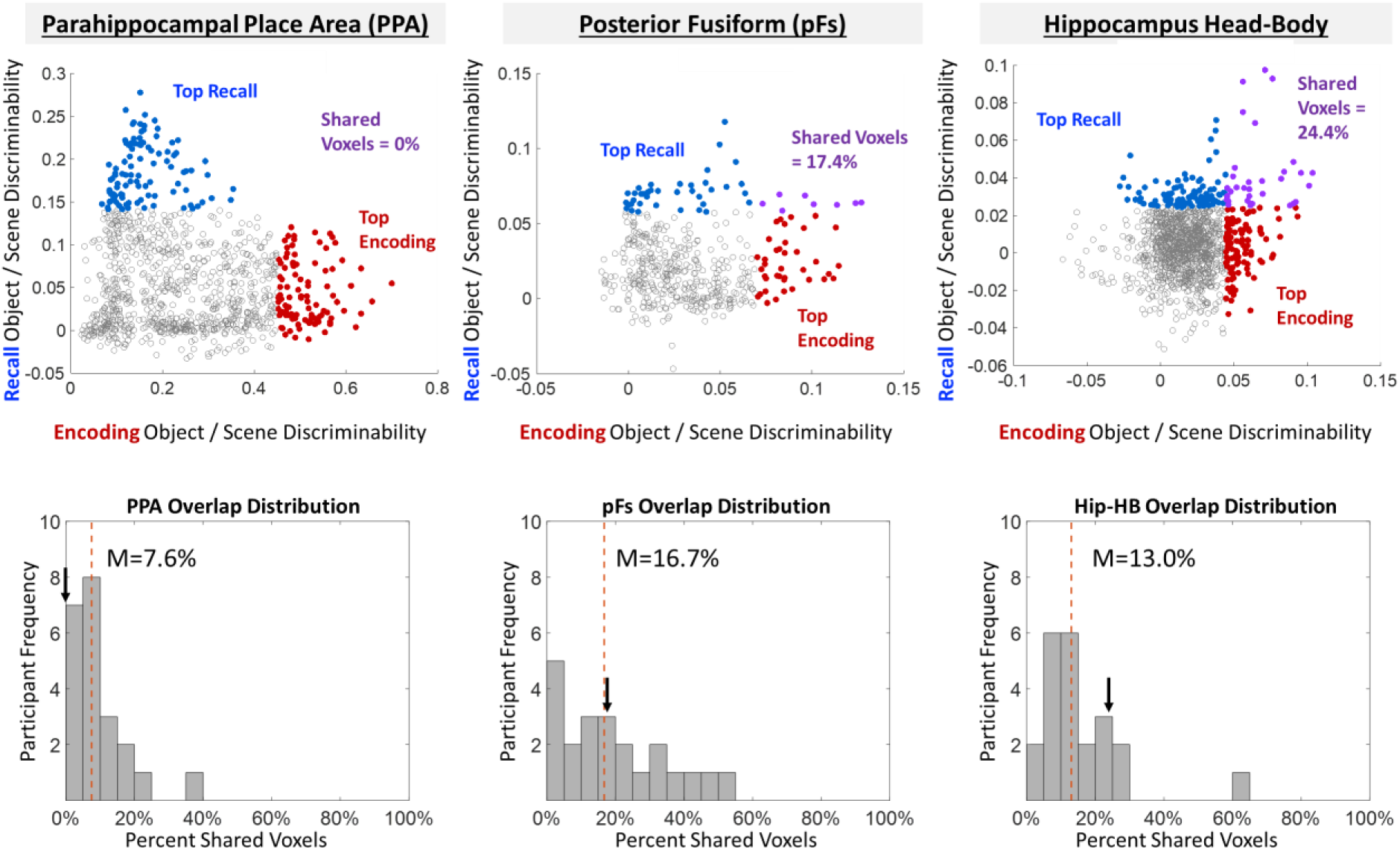
Comparing encoding and recall discriminability within the ROIs. (Top) Example ROIs from a single participant, where each point represents a voxel-centered spherical searchlight in that ROI and is plotted by the object/scene discrimination index during encoding (x-axis) versus the object/scene discrimination index during recall (y-axis). The 10% of searchlights showing strongest recall discriminability are colored in blue, while the 10% of searchlights showing strongest encoding discriminability are colored in red. Searchlights that overlap between the two (those that demonstrate both encoding and recall discrimination) are colored in purple. The patterns in this participant mirror the patterns found across participants—PPA shows low (in this case no) overlap, while pFs shows higher overlap. (Bottom) Histograms for these ROIs showing participant distribution of the percentage of overlap between the top 10% of encoding discriminating and top 10% of recall discriminating voxels. The arrow represents the participant’s data plotted above, while the dashed red line shows the median overlap percentage across participants.

To compare encoding and recall discriminability further, we focused on the top 10% of searchlights that showed encoding discriminability and compared their overlap with the top 10% of searchlights that showed recall discriminability within each ROI (Figure 6, Methods). If the same voxels perform encoding and recall discrimination, we should find significantly higher overlap than chance (approaching 100%). Conversely, if encoding and recall information comprise distinct sets of voxels, we should find equal or lower overlap compared to chance (∼10%, estimated by 100 permuted shuffles). PPA showed significantly lower overlap than chance (Median = 9.15%, Wilcoxon rank sum test: *Z* = 1.96, *p* = 0.050), while pFs showed significantly higher overlap (M = 19.14%, *Z* = 2.05, *p* = 0.040). All other regions showed no significantly different overlap than predicted by chance (MPA: 16.9%; OPA: 11.2%; LO: 14.03%; Hip-HB: 15.6%; Hip-T: 15.4%; PRC: 13.3%; PHC: 20.2%; all *p* > 0.10). These results suggest a limited relationship between encoding and recall across all visual and memory regions. pFs shows high overlap and significant correlation between encoding and recall searchlights, in addition to significant cross-discrimination across encoding and recall, suggesting some shared neural substrate. In contrast, PPA shows no correlation and significantly low overlap in addition to an absence of cross-discrimination, suggesting distinct neural substrates between encoding and recall. The remaining ROIs show mixed evidence, with relatively low correlations between encoding and recall and no difference in overlap from chance, suggesting limited shared information between encoding and recall.

#### Whole-Brain Investigation of Encoding and Recall Effects

Given the differences we observed between encoding and recall within ROIs, we conducted follow-up analyses at the whole-brain level. Looking at a group univariate contrast of objects versus scenes during encoding (Figure 7), we confirm that stimulus class selectivity is strongest in ROIs predicted by the literature: LO and pFs show high sensitivity to objects, while PPA, MPA, and OPA show high sensitivity to scenes (e.g., Epstein and Kanwisher 1998; Grill-Spector et al. 2001). Interestingly, in EVC we observe stronger responses for scenes during encoding and stronger responses for objects during recall. This may explain the negatively trending cross-discrimination between encoding and recall in EVC. However, a group univariate contrast of objects versus scenes during recall reveals that recall scene-selectivity appears strongest in areas anterior to PPA and MPA, and recall object-selectivity appears strongest in areas anterior to LO and in early visual cortex. We quantified this observation by comparing the locations of the peak encoding voxel and the peak recall voxel around each ROI for every participant. Recall peaks were significantly anterior to encoding peaks across participants bilaterally in PPA (Left Hemisphere: Wilcoxon signed rank test *Z* = 2.32, *p* = 0.020; Right Hemisphere: *Z* = 2.65, *p* = 0.008) and OPA (LH: *Z* = 2.52, *p* = 0.012; RH: *Z* = 2.91, *p* = 0.004), and in the left pFs (*Z* = 2.68, *p* = 0.007), left MPA (*Z* = 2.45, *p* = 0.014), and right LO (*Z* = 2.71, *p* = 0.007). Even in those hemispheres showing non-significant effects the same numeric trend was observed, with recall peaks anterior to encoding peaks. In sum, rather than the peaks of recalled stimulus class overlapping with those of encoding, the greatest scene-object differences occur in a spatially separate set of voxels largely anterior to those during encoding.

**Figure 7.**
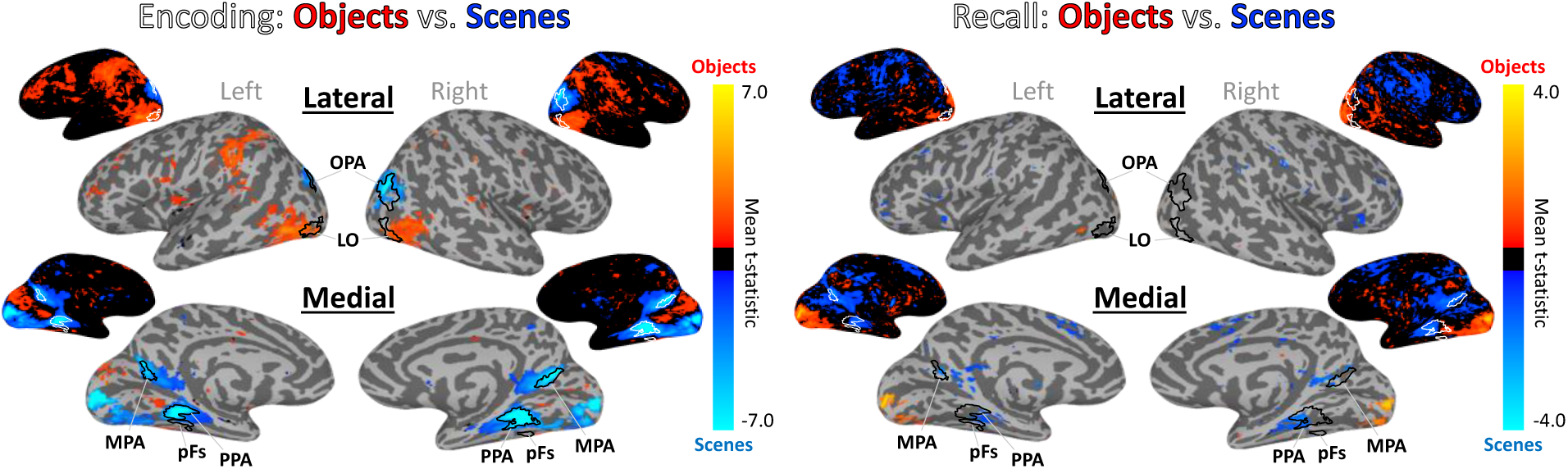
Whole-brain activation of objects and scenes during encoding and recall. Univariate whole-brain t-statistic maps of the contrast of objects (red/yellow) versus scenes (blue/cyan) in encoding (left) and recall (right). Contrasts show group surface-aligned data (N=22), presented on the SUMA 141-subject standard surface brain (Saad and Reynolds 2012). Outlined ROIs are defined by voxels shared by at least 25% participants from their individual ROI definitions (using independent functional localizers), with the exception of the pFs and OPA which were defined by 13% overlap (there were no voxels shared by 25% of participants). The encoding maps are thresholded at FDR corrected *q*<0.05. For the recall maps, no voxels passed FDR correction, so the contrast presented is thresholded at *p*<0.01 for visualization purposes. Smaller surface maps show unthresholded results.

A searchlight analysis looking at information discriminability across the brain replicates this spatial separation (Figure 8). During encoding, scenes and objects are most discriminable in the same regions identified by the independent perceptual localizer (LO, pFS, PPA, MPA, OPA). However, during recall, peak discriminability visibly occurs in voxels anterior to these encoding-based regions. A comparison of the peak voxel locations between encoding and recall confirmed that recall was significantly anterior to encoding in several regions (right PPA: *Z* = 2.06, *p* = 0.039; left OPA: *Z* = 3.20, *p* = 0.001; left LO: *Z* = 2.61, *p* = 0.009; right LO: *Z* = 2.06, *p* = 0.039; left pFs: *Z* = 2.21, *p* = 0.027), and numerically showing the same trend in others (left PPA, left and right MPA, right OPA). Next, we employed a cross-discrimination searchlight to identify regions with shared stimulus representations between encoding and recall. Again, areas anterior to those most sensitive during encoding showed highest similarity between encoding and recall representations. This anterior shift was significant in bilateral PPA (LH: *Z* = 3.30, *p* = 9.83 × 10^-4^; RH: *Z* = 2.39, *p* = 0.017), bilateral OPA (LH: *Z* = 3.59, *p* = 3.34 × 10^-4^; RH: *Z* = 3.43, *p* = 6.15 × 10^-4^), left pFs (*Z* = 2.38, *p* = 0.017), right LO (*Z* = 2.97, *p* = 0.003) and numerically showed the same trend in left LO, right pFs, and right MPA.

**Figure 8.**
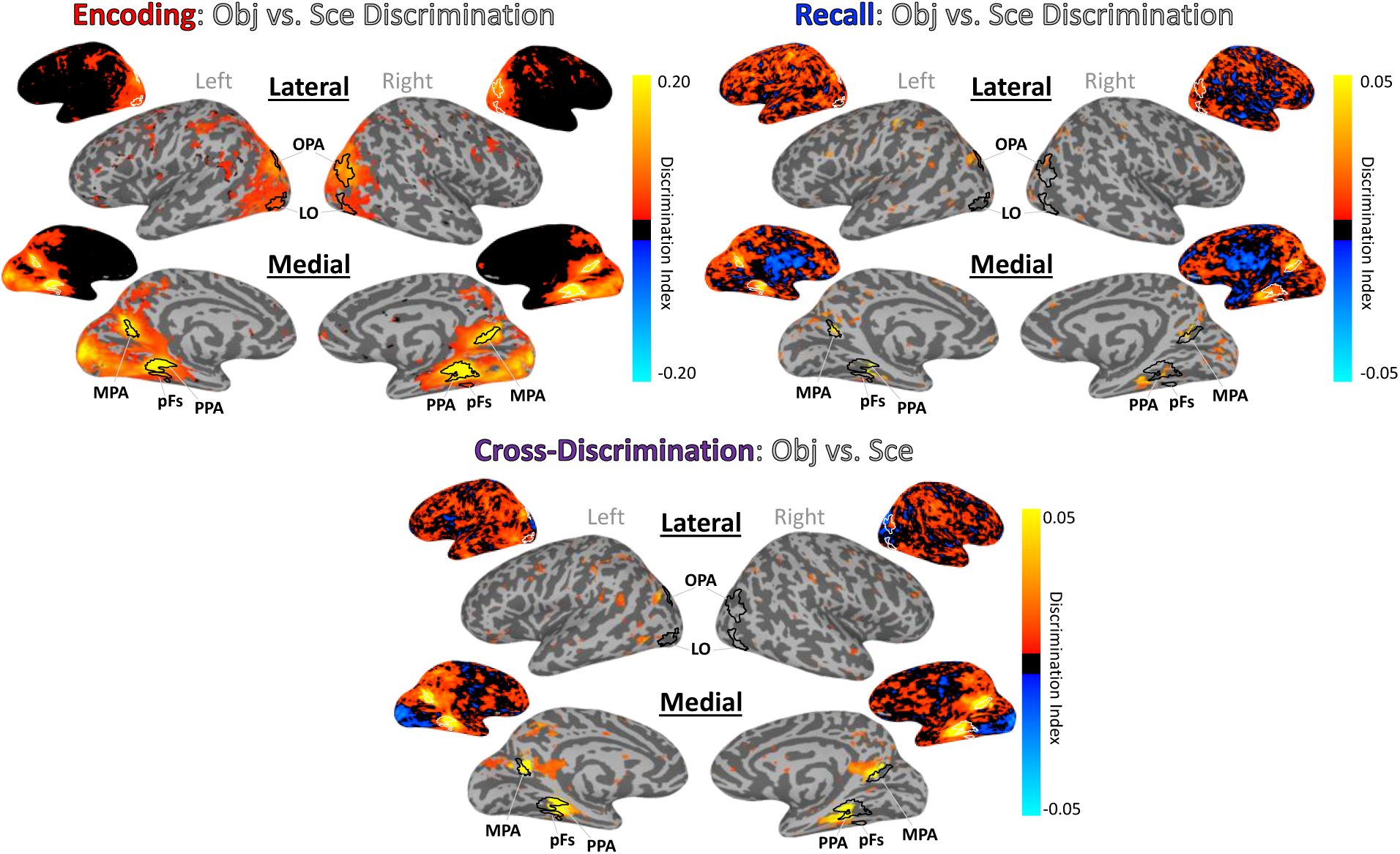
Whole-brain discrimination analyses for encoding, recall, and cross-decoding of information. Whole-brain searchlight analyses investigating discrimination of objects versus scenes during encoding (top left), recall (top right), and cross-discrimination (bottom). Brighter yellows indicate higher discrimination indices. Outlined ROIs are defined using independent stimuli in an independent localizer run. All maps are thresholded at *p*<0.005 uncorrected, and unthresholded maps are also shown. The cross-discrimination searchlight shows regions that have a shared representation between encoding and recall.

These results suggest a spatial separation between encoding and recall with strongest reinstatement occurring outside of scene- and object-selective regions typically localized in visual tasks.

## Discussion

In this work, we conducted an in-depth investigation of how and where recalled memory content for complex object and scene images is represented in the brain. First, we observed a striking difference in the representational structure between encoding and recall. While information in cortex during encoding reflected multiple levels of information, during recall we observed clear evidence for coarse-level information (objects versus scenes) as well as some fine-level scene information. No region showed mid-level information during recall (e.g., natural/manmade for scenes, tool/non-tools for objects), even though such information was often stronger than fine-level information during encoding. In hippocampus, we only observed coarse-level discrimination and only during encoding. Medial temporal lobe regions perirhinal cortex and parahippocampal cortex also only showed coarse-level discrimination, although this information was discriminable during both the encoding and recall periods. Second, a direct comparison between encoding and recall discriminability within ROIs found only weak correlations that were significant in a limited number of ROIs. When we further examined just the top discriminating voxels for encoding and recall, most regions showed no overlap between them, with only pFs showing higher overlap than chance. Finally, a whole brain comparison of encoding and recall discriminability revealed that the peaks for recall as well as the strongest encoding-recall similarity were spatially anterior to the peaks during encoding. Collectively, our results reveal key spatial and representational differences between encoding and recalling stimulus content.

The ability to decode scenes versus objects during recall is consistent with several findings showing broad stimulus class decodability during recall (Polyn et al. 2005; Reddy et al. 2010; Boccia et al. 2019; O’Craven and Kanwisher 2000). Similarly, the ability to decode fine-level information of individual scene categories is consistent with prior work showing decoding of specific stimulus images (e.g., Dickerson et al. 2007; Buchsbaum et al. 2012; Lee et al. 2012; Kuhl and Chun 2014). Additionally, we replicate several findings observing discriminability of different levels of information during perception (e.g., Mahon et al. 2007; Walther et al. 2009; Kravitz et al. 2011; Park et al. 2011). We did not find discriminability of object size in visual areas (Konkle et al. 2012) as expected, but this may reflect the range of sizes we selected, which were not as far apart as in prior work. We also find a significant ability to decode memory vividness and future recognition success from many cortical regions as shown in prior work (Supplementary Material S1, S2; Brewer et al. 1998; Wais 2008; Dijkstra et al. 2017; Fulford et al. 2018). However, at face-value the limited decoding we find during recall as well as the low encoding-recall similarity in category-selective cortex appear to be at odds with prior findings. We discuss each of these issues in turn in the paragraphs below.

While we were able to discriminate coarse-level information in most areas and fine-level information in some areas during recall, we found no evidence for recall of mid-level information in any region. Prior work has primarily focused on these coarse- and fine-levels, and this absence of mid-level information suggests that imagery-based representations in cortex do not contain more information that generalizes across categories. Participants may be recalling limited image features, sufficient for fine-level classification of some specific image categories (e.g., retinotopic features shared across exemplars of a category), and sufficient for classification at the coarse level of scenes versus objects (given large differences between their features). However, the representations during recall may not contain more abstract information, such as features shared by items at a similar mid-level (e.g., size, function, qualities of a scene). This pattern of results is reflected not only in many category-selective areas, but also in early visual cortex, which is unlikely to represent these more abstract features.

In terms of encoding-recall similarity, our results also appear to be inconsistent with some previous findings of sensory reinstatement, in which the neurons or voxels sensitive during encoding have been reported to show the same patterns during recall (Wheeler et al. 2000; Danker and Anderson 2010; Buchsbaum et al. 2012; Johnson and Johnson 2014; Tompary et al. 2016; Schultz et al. 2019). In several visual and memory-related regions, we observed limited overlap between the sub-regions with peak encoding and those with peak recall information, with the strongest encoding-recall similarity in more anterior regions. However, some studies do report encoding-recall similarity within scene- and object-selective cortex (e.g., O’Craven and Kanwisher 2000; Johnson and Johnson 2014) which may be attributable to key methodological differences from the current study. First, as noted above, we targeted recollection of stimulus content rather than individual items. While scene- and object-selective regions may maintain item-specific visual information during both encoding and recall, our results suggest a difference in representations during encoding and recall at more generalized levels of information. Second, we employed an item-based recall task, rather than associative tasks commonly used to study recall (e.g., Ganis et al. 2004; Zeidman et al. 2015a; Xiao et al. 2017; Jonker et al. 2018). This allowed us to ensure that information we decoded was not related to other factors such as decoding a cue or association. One potentially interesting question for future work is how strength of the memory representation (e.g., reported vividness) or task may modulate the degree of overlap between encoding and recalled representations.

Our findings suggest a posterior-anterior gradient within cortical regions, in which recalled representations extend anterior to encoding or perceptual representations. These results agree with recent research showing that regions involved in scene memory are anterior to those involved in scene perception, with the possibility of separate perception and memory networks (Baldassano et al. 2016; Burles et al. 2018; Chrastil 2018; Silson et al. 2019a). This anterior bias for recall may reflect top-down refreshing of a memory representation in contrast to the largely bottom-up processes that occur during perception (Mechelli et al. 2004; Johnson et al. 2007; Dijkstra et al. 2019). Indeed, recent work using electroencephalography (EEG) has identified a reversal of information flow during object recall as compared to encoding (Linde-Domingo et al. 2019). Alternatively, other research has suggested a gradient within the neocortex that reflects a split of conceptual information represented anterior (or downstream) to perceptual information (Peelen and Caramazza 2012; Borghesani et al. 2016; Martin 2016). While recent work shows highly detailed visual content within recalled memories (Bainbridge et al. 2019), it is possible recalled memories may be more abstracted and conceptual compared to their encoded representations. This recalled memory could thus contain less mid-level perceptual information or be abstracted into a different representation, explaining why we can decode memory strength but not fine-grained perceptually-defined distinctions (e.g., natural versus manmade) during recall. Collectively, our results support these two possible accounts for anterior-posterior gradients of memory/perception or conceptual/perceptual information in the brain, in contrast with other accounts claiming an identical representation between encoding and recall (e.g., Schultz et al. 2019).

The current work also provides further support for a content-independent role of the hippocampus in memory. During encoding, we observe broad content selectivity in the hippocampus, as has been observed in other recent work claiming a perceptual role for the hippocampus (Zeidman et al. 2015b; Hodgetts et al. 2017). However, we do not observe strong evidence of any other content representations during encoding or recall; the hippocampus does not show sensitivity to more fine-grained information, and during recall, it does not even show differences at the broadest distinction of objects versus scenes. These results lend support for the notion that the hippocampus is largely content-independent (Davachi 2006; Danker and Anderson 2010; Liang et al. 2013; Schultz et al. 2019), with individual item decoding in previous work possibly driven by decoding of indexes within the hippocampus connected to fuller representations in the neocortex, or a coding of memory strength (e.g., Teyler and Rudy, Jonker et al. 2018). In fact, while stimulus content during recall is not discriminable, we find that memory strength is decodable from the hippocampus, mirroring similar results finding strength but not content representations in the hippocampus for oriented gratings (Bosch et al. 2014).

There is also evidence to suggest that the hippocampus may require longer delays (e.g., several hours to a week) to develop decodable representations of memory content (Tompary and Davachi 2017; Lee et al. 2019), and so a similar experiment conducted with longer delays between study and test (e.g., days) may find decodable stimulus content from the hippocampus. These results in the hippocampus also serve as an interesting counterpoint to our findings in PRC and PHC within the medial temporal lobe. While these regions also do not show any mid- or fine-grained information, they show significant discrimination of coarse-level information during encoding, recall, and cross-discriminable representations between the two. These results support prior findings of category selectivity within these regions (e.g., Murray & Richmond 2001; Buckley & Gaffan 2006; Staresina et al. 2011) as well as work suggesting similar representations between perception and recall (Schultz et al. 2019).

The current study combining nested categorical structure for real-world images and an item-based recall approach allows us to observe different levels of stimulus representations across the brain; however, there are limitations to this methodology that could be addressed in future work. In particular, the limited information and null findings during recall could partly reflect a lack of power reflecting the weaker signals during recall compared to encoding. However, from a combination of our results, we think issues of power alone cannot explain our findings. First, several regions during encoding show stronger mid-level discriminability than fine-level discriminability (e.g., PPA, pFs, and MPA for natural/manmade). However, these same regions show significant fine-level discriminability but not mid-level discriminability during recall, suggesting that the nature of the information present during recall is different, not just diminished. Second, our ability to decode recall vividness from most visual regions suggests decodable patterns of information are present during recall (Supplementary Material S1, S2). Third, our analyses uncovering separate peaks of encoding and recall also suggest that significant recall discriminability does exist, but in regions somewhat distinct from these perceptually-based ROIs. Finally, the current sample size (N=22) and number of trials (192 stimuli) fall in the higher range compared to related studies (e.g., Lee et al. 2012: N=11; Johnson and Johnson 2014: N=16; Schultz et al. 2019: N=16). Another limitation of our current study is our inability to assess discriminability for individual images – our current methodology was designed to allow us to powerfully test stimulus content divorced from memory for individual items. Future studies should investigate whether individual item representations are identical between encoding and recall, even if more general content representations are not. Such findings could have meaningful implications on the nature of representations during recall, suggesting the imagery for an individual item is vivid enough to be item-specific, but results in a limited level of abstraction. Another question for future research will be a deeper examination of the different factors influencing encoding-recall similarity for each ROI. While the conjunction of our results in addition to the whole-brain analyses suggests a clear difference between encoding and recall, high encoding-recall correlations or low overlap in isolation could be attributed to alternate explanations. For example, high correlations between encoding and recall could be due to anatomical influences on voxel activity. On the other hand, at-chance overlap between encoding and recall could be due to high noise within an ROI. Finally, it will be important to see whether these newly defined anterior recall regions show more fine-grained representations of stimulus content during recall, and whether there may be region-specific differences (e.g., the MPA in particular has been a key target for comparisons of scene perception and scene recall; Burles et al. 2018; Chrastil 2018; Silson et al. 2019b).

Examining item-based recall and representations of memory content in the brain has ultimately unveiled a rather complex, nuanced relationship of encoding and recall, with strongest encoding-recall similarity occurring largely anterior to scene- and object-selective visual cortex. In the study of memory, it is important to examine not only how we remember, but *what* we are remembering, and this study reveals that the way in which this content is manifested may vary greatly between encoding and recall.

## Acknowledgements

We thank Adam Steel, Caroline Robertson, and Alexa Tompary for their helpful comments on the manuscript. This research was supported by the Intramural Research Program of the National Institutes of Health (ZIA-MH-002909), under National Institute of Mental Health Clinical Study Protocol 93-M-1070 (NCT00001360).

