## Supplementary Material for "Distinct representational structure and localization for visual encoding and recall during visual imagery"

**Title:** Distinct representational structure and spatial distribution for visual encoding and recall

**Abbreviated title:** Distinct representations for encoding and recall

**Table of Contents**

pages
3-6	**Supplementary Material 1:** Classification of behavior

7	**Supplementary Material 2:** *Figure SF1.* Relationship of fMRI response patterns to

behavior

8	**Supplementary Material 3:** *Figure SF2.* Information discriminability in

hippocampal subfields

9	**Supplementary Material 4:** *Table ST1.* ROI voxel sizes by participant

10	**Supplementary Material 5:** *Table ST2.* Permutation-based significance for

discriminability analyses

11	**Supplementary Material 6:** *Table ST3.* Discrimination indices for all ROIs.

12-14	**Supplementary Material 7:** ROI-to-ROI similarity analysis

15	**Supplementary Material 8:** *Figure SF3.* Comparison of stimulus representations

between regions

16	**Supplementary References**

Supplementary Material 1: Classification of behavior

**Methods**

We conducted analyses to relate fMRI activity to behavioral measures of reported recall vividness and post-scan recognition performance. An additional general linear model (GLM) was conducted to obtain an estimate for each trial for each run, so that they could be compared with trial-by-trial behavior measures. In this case, no memory trials were retained in the GLM, so that we could look at ability to classify memory vividness responses.

To perform classification on recall vividness, we employed support vector machines (SVMs) to classify recall vividness responses (no memory, low vividness, high vividness) from ROI patterns during encoding or recall. These SVMs were conducted on all pairs of the three classes (high vs. low, low vs. no, and high vs. no), and then classification accuracies were averaged to give a combined classification metric. These classification accuracies were compared against a chance classification level of 50%. We also report the results of classifying low versus high vividness trials, without no memory trials. Classifications were conducted using a leave-one-run-out approach (i.e., training on 7 runs of data and testing on 1 held-out run, across all 8 possible splits of the runs). For all classifications, training and testing sets were first downsampled to create equally sized classes. A random number of datapoints was taken from each class to match the size of the smallest set. The minimum number of trials needed for a classification to be conducted within a participant was 16 trials per class (14 training, 2 testing). All participants passed this criterion for high vividness trials, while 21 out of 22 passed for low vividness trials, and 7 passed for no memory trials. Thus note that the 3-class average classification accuracy is based on the average across a small number of participants (N=7), while the high/low classification accuracy is an average across 21 participants.

We also used SVMs to measure the ability to classify later successful recognition versus failed recognition (hits vs. misses) from ROI patterns during encoding or recall. These classification accuracies were compared against a chance classification level of 50%. Again, all SVMs were conducted using a leave-one-run-out approach. Within each participant, training and testing sets were downsampled so that both classes (hits and misses) had equal sizes, matching the size of the smallest class. The minimum number of trials per participants required for the classification to be performed was 16 trials per class (14 for training, 2 for testing). All participants passed this criterion. We focused these analyses on ROIs in category-selective cortex and hippocampus.

**Results and Discussion**

*FMRI response patterns and memory vividness*

On average, participants reported ‘high vividness’ on 60.8% of trials (SD=16.9%), ‘low vividness’ on 29.9% of trials (SD=12.8%), and ‘no memory’ on 9.31% of trials (SD=8.38%). We tested the ability to classify this memory vividness from patterns in the different ROIs during encoding and recall (Figure SF1). Encoding patterns from all scene- and object-selective regions were predictive of recall vividness: LO (Mean classification accuracy = 62.28%, *p =* 3.09 × 10^-5^), pFs (*M* = 60.90%, *p =* 9.99 × 10^-5^), PPA (*M* = 57.11%, *p =* 8.12 × 10^-5^), MPA (*M* = 55.72%, *p =* 0.027), and OPA (*M* = 57.51, *p =* 4.20 × 10^-4^). In contrast, within the hippocampus, (Hip-HB and Hip-T), encoding patterns could not be used to classify recall vividness (both *p* > 0.05). Recall patterns from all scene- and object-selective regions were also predictive of recall vividness: LO (*M* = 56.43%, *p =* 4.61 × 10^-4^), pFs (*M* = 55.83%, *p =* 0.003), PPA (*M* = 55.63%, *p =* 0.016), MPA (*M* = 57.50%, *p =* 0.007), and OPA (*M* = 58.26%, *p =* 0.0012). Additionally, the hippocampus showed memory vividness decodability from patterns during recall, in both the Hip-HB (*M* = 53.34%, *p =* 0.012) and Hip-T (*M* = 54.26%, *p =* 0.022). While hippocampus showed decodability during recall and not during encoding, there was no significant difference in classification accuracy between encoding and recall (both regions: *p* > 0.05).

If we look specifically at the ability to classify low vividness from high vividness trials, a subset of these ROIs shows above-chance classification accuracy. Specifically, during memory encoding, patterns in the pFs (*M* = 62.35%, *p* = 0.013) and OPA (*M* = 58.70%, *p* = 2.33 × 10^-4^) can differentiate low and high vividness trials. During recall, pFs (*M* = 57.22%, *p* = 0.048), PPA (*M* = 61.07%, *p* = 0.012), and OPA (*M* = 59.20%, *p* = 0.046) patterns can differentiate low vividness from high vividness trials. The hippocampus did not show ability to differentiate low vividness or high vividness trials during encoding nor recall (both regions: *p* > 0.10).

These results indicate that scene- and object-selective regions contain information about memory vividness during both encoding and recall. In contrast, the hippocampus only contains information about memory vividness during the recall period, when classifying all trial types (high vivid, low vivid, and no memory). However, it is less clear whether the hippocampus shows decoding of more fine-grained differences in vividness (differentiating high versus low vividness).

*FMRI response patterns and subsequent memory*

Encoding patterns from scene- and object-selective regions were predictive of later successful recognition (Figure SF1), from the LO (*M* = 53.08%, *p* = 0.032), PPA (*M* = 52.08%, *p* = 0.033), and OPA (*M* = 53.35%, *p* = 0.001). MPA, pFs, Hip HB and Hip T did not show significant decodability of successful recognition (all *p* > 0.10). No recall patterns from any ROI were predictive of later successful recognition (all *p* > 0.10). These results confirm prior findings demonstrating an ability to predict subsequent memory performance from visual areas (Brewer et al. 1998), and show little ability to decode later recognition success from neural information during the recall period.

Supplementary Material 2

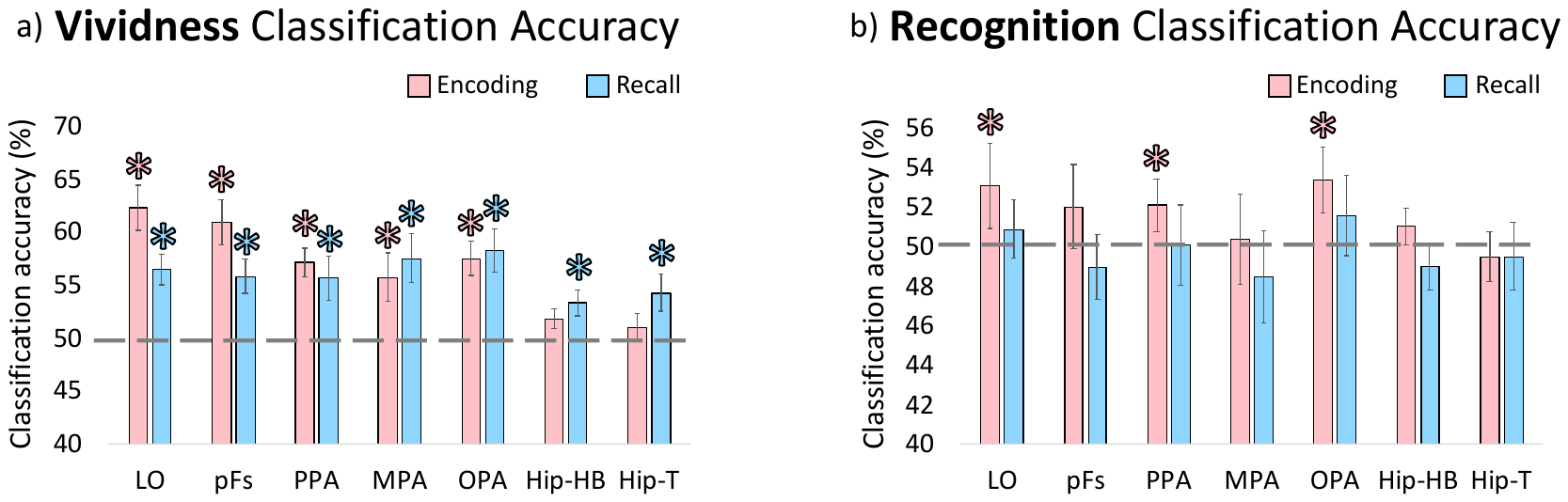

**Figure SF1** – **Relationship of fMRI response patterns to behavior.** (a) Classification accuracies for predicting subjectively reported recall vividness, based on patterns of encoding (pink) and recall (blue) from the ROIs. Chance classification level is 50%, and marked with the dashed grey line. Asterisks indicate a significantly higher classification accuracy than chance (*p* < 0.05). In scene- and object-selective visual regions, patterns during both encoding and recall can significantly predict the vividness of recall. In the hippocampus, only patterns during recall can significantly predict memory vividness. (b) Classification accuracies for predicting post-scan memory performance in a recognition task from patterns of encoding (pink) and recall (blue) from the ROIs. Only patterns from some scene- and object-selective regions during encoding (LO, PPA, OPA) show significant ability to predict later recognition success. No patterns during recall were linked with recognition success.

Supplementary Material 3

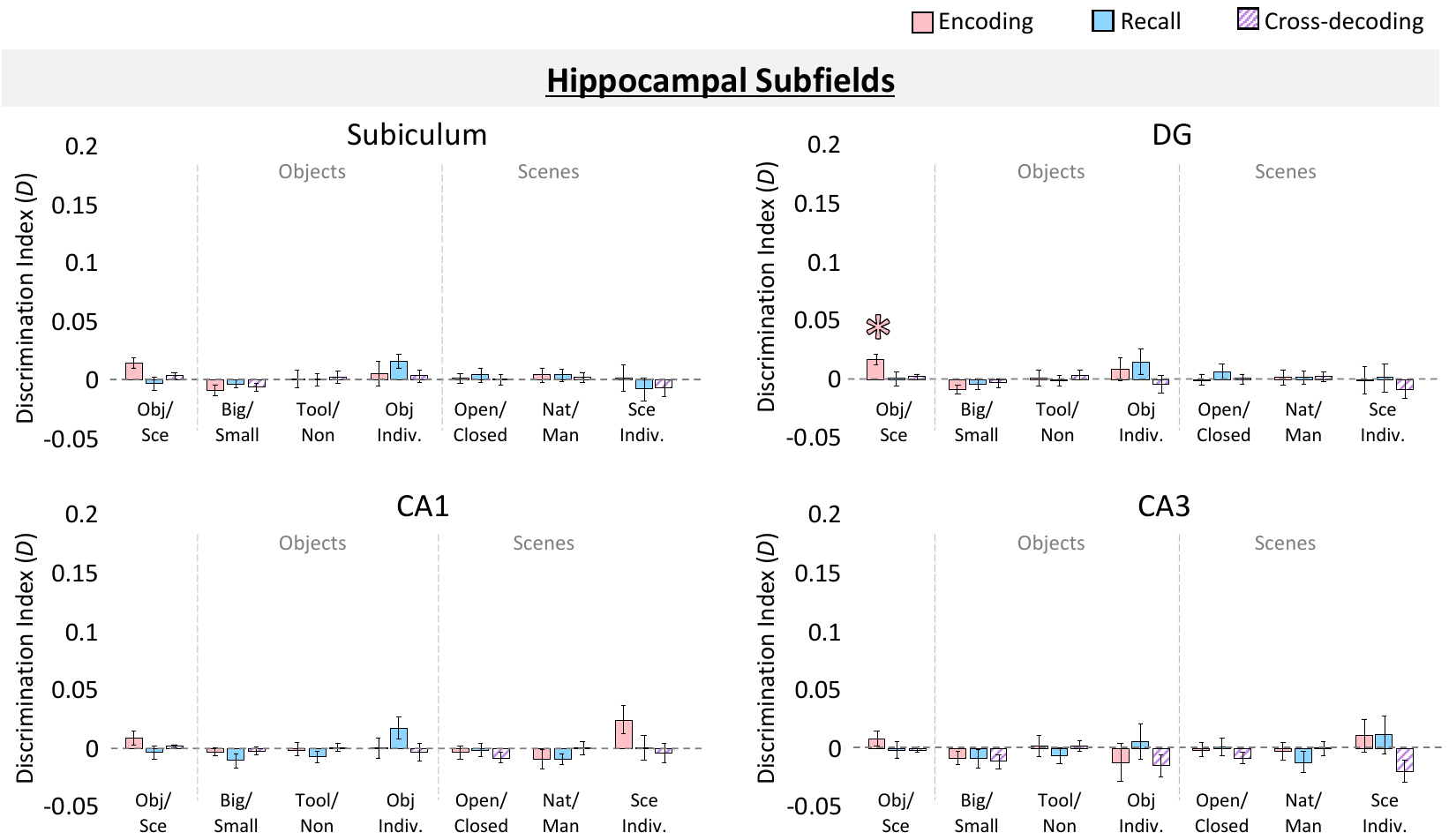

**Figure SF2 – Information discriminability in hippocampal subfields**. Bar graphs are displayed in the same manner as Figure 4 and 5 in the main manuscript, indicating mean discrimination index for comparisons of different levels of stimulus information (class, type, and category for objects and scenes) for different types of memory trials (encoding, recall, and cross-decoding between encoding and recall). Shown are discriminability indices for the hippocampal subfields subiculum, dentate gyrus (DG), *Cornu Ammonis* 1 (CA1), and *Cornu Ammonis* 3 (CA3). Subfields were automatically segmented using FreeSurfer’s hippocampal subfield segmentation tool. Error bars indicate standard error of the mean. Asterisks (*) indicate significance at a FDR corrected level of *q* < 0.05.

Supplementary Material 4

Table ST1. **ROI voxel sizes by participant**. This table shows the number of voxels contained within each region of interest (ROI) for each of the 22 participants. For the functionally defined regions, “L” indicates the left hemisphere ROI, while “R” indicates the right hemisphere ROI. A dash indicates that that ROI was not identified in that participant. Both hemispheres were combined to make a bilateral ROI for all analyses.

|  | **L-LO** | **R-LO** | **L-pfS** | **R-pFs** | **L-PPA** | **R-PPA** | **L-MPA** | **R-MPA** | **L-OPA** | **R-OPA** | **Hip-HB** | **Hip-T** | **PRC** | **PHC** | **EVC** |
| --- | --- | --- | --- | --- | --- | --- | --- | --- | --- | --- | --- | --- | --- | --- | --- |
| **1** | 344 | 551 | 472 | 118 | 567 | 446 | 347 | 718 | 366 | 591 | 2067 | 579 | 876 | 2493 | 2155 |
| **2** | 1086 | 937 | 231 | 302 | 259 | 418 | 268 | 356 | 50 | 48 | 2486 | 741 | 793 | 2250 | 3386 |
| **3** | 1322 | 680 | 121 | 129 | 1087 | 1642 | 934 | 1265 | 1520 | 1962 | 1918 | 645 | 729 | 1083 | 4289 |
| **4** | 84 | 307 | 1188 | 506 | 40 | 2000 | 30 | - | - | 90 | 2675 | 751 | 827 | 1381 | 1700 |
| **5** | 544 | 538 | 886 | 582 | 791 | 674 | 535 | 1529 | 488 | 759 | 2291 | 630 | 1073 | 1828 | 2580 |
| **6** | 115 | 358 | 214 | 301 | 161 | 495 | - | - | - | - | 2498 | 570 | 926 | 2200 | 915 |
| **7** | 94 | 193 | 58 | 102 | 23 | 85 | - | - | 157 | 144 | 2795 | 706 | 1924 | 2782 | 4972 |
| **8** | 1218 | 917 | 406 | 655 | 176 | 256 | 92 | 108 | 207 | 257 | 2184 | 548 | 709 | 1998 | 1562 |
| **9** | 1910 | 1065 | 469 | 416 | 550 | 511 | 99 | 337 | 824 | 1282 | 1757 | 461 | 804 | 1720 | 3414 |
| **10** | 2691 | 1379 | 238 | 404 | 544 | 801 | 405 | 620 | 138 | 232 | 2684 | 716 | 946 | 1995 | 1144 |
| **11** | 1056 | 841 | 500 | 309 | 639 | 1222 | 400 | 706 | 505 | - | 2551 | 695 | 1392 | 2085 | 6017 |
| **12** | 630 | 1133 | 226 | 231 | 376 | 620 | 1547 | 732 | 584 | 256 | 1308 | 346 | 1120 | 2479 | 2303 |
| **13** | 1610 | 1738 | 1005 | 542 | 235 | 596 | - | 1875 | 97 | 106 | 2261 | 747 | 1298 | 2539 | 4186 |
| **14** | 244 | 515 | 126 | 136 | 289 | 638 | 111 | 407 | 75 | 798 | 1937 | 309 | 993 | 2632 | 1075 |
| **15** | 3354 | 635 | 245 | 76 | 640 | 966 | 2338 | 899 | 1529 | 1415 | 2382 | 736 | 936 | 2103 | 2940 |
| **16** | 1527 | 2394 | 511 | 461 | 95 | 259 | 48 | 50 | 24 | 88 | 2215 | 594 | 839 | 1681 | 2205 |
| **17** | 1778 | 165 | 294 | 270 | 143 | 512 | - | 110 | 101 | 693 | 1944 | 279 | 1139 | 2059 | 2342 |
| **18** | 1405 | 552 | 449 | 141 | 325 | 598 | 110 | 197 | 565 | 1110 | 2444 | 657 | 1091 | 2252 | 1616 |
| **19** | 815 | 1264 | 294 | 291 | 453 | 597 | 114 | 207 | 2853 | 795 | 2143 | 695 | 896 | 2421 | 1930 |
| **20** | 391 | 84 | 147 | 60 | 622 | 520 | 402 | 492 | 714 | 134 | 2280 | 701 | 1002 | 2562 | 3118 |
| **21** | 1267 | 1251 | 436 | 322 | 285 | 347 | 192 | 459 | 71 | 429 | 2990 | 898 | 918 | 2321 | 6549 |
| **22** | 683 | 380 | 348 | 467 | 63 | 262 | 167 | 161 | 42 | 22 | 3056 | 850 | 1047 | 2406 | 6551 |
| **MEAN** | 1098.5 | 812.6 | 402.9 | 310.0 | 380.1 | 657.5 | 452.2 | 590.9 | 545.5 | 560.6 | 2312.1 | 629.7 | 1012.6 | 2148.6 | 3043.1 |
| **SD** | 844.1 | 560.6 | 289.6 | 176.0 | 274.7 | 453.0 | 600.9 | 503.0 | 704.7 | 540.7 | 409.8 | 161.7 | 265.7 | 418.0 | 1722.7 |

Supplementary Material 5

Table ST2. **Permutation-based significance for discriminability analyses**. *P*-values in the discriminability analysis when using permutation tests. Permutations were conducted as 1,000 randomizations of each RSM per participant, from which permuted discrimination indices were calculated. True and permuted discrimination indices were compared across participants with one-tailed t-tests. The p-threshold indicates the FDR cutoff for significance, and significant discriminations are colored based on the key.

| **Scenes /**  **Objects** | | Enc | Rec | Cross | *p-threshold* | | |  | | |  |  |  |  |  |  |
| --- | --- | --- | --- | --- | --- | --- | --- | --- | --- | --- | --- | --- | --- | --- | --- | --- |
| Obj. ROIs | LO | 1.0E-5 | 0.001 | 0.23 |  | 0.009 | |  | | |  | | |  | | |
|  | pFs | 3.1E-5 | 0.005 | 0.006 |  | 0.011 | |  | | |  | | | Significant Encoding Discriminability (FDR-corrected) | | |
| Sce. ROIs | PPA | 3.3E-7 | 3.4E-4 | 0.034 |  | 0.008 | |  | | |  | | | Significant Recall Discriminability (FDR-corrected) | | |
|  | MPA | 3.1E-6 | 0.006 | 1.5E-4 |  | 0.007 | |  | | |  | | | Significant Cross Discriminability (FDR-corrected) | | |
|  | OPA | 1.3E-6 | 0.004 | 0.67 |  | 0.004 | |  | | |  | | | *p* < 0.05 (does not pass FDR correction) | | |
| Hipp. | HB | 2.9E-4 | 0.81 | 0.08 |  | 2.9E-4 | |  | | |  | | |  | | |
|  | T | 0.11 | 0.48 | 0.11 |  | None | |  | | |  | | |  | | |
| Other | PRC | 5.8E-5 | 0.004 | 2.5E-4 |  | 0.004 | |  | | |  | | |  | | |
|  | PHC | 4.4E-6 | 0.003 | 4.1E-4 |  | 0.003 | |  | | |  | | |  | | |
|  | EVC | 4.4E-7 | 0.004 | 0.99 |  | 0.007 | |  | | |  | | |  | | |
| **Objects** | | Big vs. Small | | | Tools vs. Non-tools | | | | Object Individuation | | | | | | | |
|  |  | Enc | Rec | Cross | Enc | Rec | Cross | | Enc | | | Rec | | | Cross | |
| Obj. ROIs | LO | 0.16 | 0.58 | 0.71 | 0.005 | 0.13 | 0.28 | | 1.5E-4 | | | 0.71 | | | 0.18 | |
|  | pFs | 0.12 | 0.93 | 0.90 | 0.011 | 0.92 | 0.47 | | 1.3E-4 | | | 0.12 | | | 0.18 | |
| Sce. ROIs | PPA | 0.53 | 0.88 | 0.63 | 0.11 | 0.96 | 0.46 | | 2.2E-4 | | | 0.09 | | | 0.62 | |
|  | MPA | 0.73 | 0.99 | 0.58 | 0.84 | 0.96 | 0.79 | | 0.049 | | | 0.09 | | | 0.053 | |
|  | OPA | 0.72 | 0.89 | 0.70 | 0.76 | 0.95 | 0.58 | | 7.5E-5 | | | 0.17 | | | 0.13 | |
| Hipp. | HB | 0.97 | 0.95 | 0.92 | 0.38 | 0.84 | 0.27 | | 0.32 | | | 0.009 | | | 0.63 | |
|  | T | 0.91 | 0.87 | 0.89 | 0.69 | 0.35 | 0.51 | | 0.54 | | | 0.40 | | | 0.40 | |
| Other | PRC | 0.78 | 0.74 | 0.84 | 0.39 | 0.74 | 0.23 | | 0.17 | | | 0.35 | | | 0.34 | |
|  | PHC | 0.66 | 0.77 | 0.39 | 0.47 | 0.75 | 0.52 | | 0.35 | | | 0.35 | | | 0.41 | |
|  | EVC | 0.12 | 0.86 | 0.97 | 0.67 | 0.37 | 0.23 | | 5.2E-7 | | | 0.007 | | | 0.99 | |
| **Scenes** | | Open vs. Closed | | | Natural vs. Manmade | | | | | Scene Individuation | | | | | | |
|  |  | Enc | Rec | Cross | Enc | Rec | Cross | | | Enc | | | Rec | | | Cross |
| Obj. ROIs | LO | 0.009 | 0.85 | 0.25 | 0.001 | 0.91 | 0.97 | | | 0.039 | | | 0.09 | | | 0.68 |
|  | pFs | 0.001 | 0.65 | 0.32 | 0.023 | 0.61 | 0.39 | | | 0.034 | | | 0.011 | | | 0.37 |
| Sce. ROIs | PPA | 0.001 | 0.76 | 0.28 | 0.002 | 0.55 | 0.033 | | | 0.001 | | | 0.008 | | | 0.46 |
|  | MPA | 0.027 | 0.39 | 0.70 | 4.3E-4 | 0.40 | 0.33 | | | 0.06 | | | 0.007 | | | 0.28 |
|  | OPA | 0.17 | 0.65 | 0.95 | 0.055 | 0.69 | 0.24 | | | 0.002 | | | 0.049 | | | 0.32 |
| Hipp. | HB | 0.44 | 0.27 | 0.76 | 0.44 | 0.67 | 0.28 | | | 0.22 | | | 0.53 | | | 0.93 |
|  | T | 0.021 | 0.054 | 0.16 | 0.14 | 0.65 | 0.39 | | | 0.97 | | | 0.24 | | | 0.86 |
| Other | PRC | 0.032 | 0.59 | 0.50 | 0.26 | 0.82 | 0.43 | | | 0.83 | | | 0.26 | | | 0.69 |
|  | PHC | 0.013 | 0.23 | 0.23 | 0.10 | 0.69 | 0.36 | | | 0.34 | | | 0.13 | | | 0.83 |
|  | EVC | 0.19 | 0.91 | 0.56 | 0.046 | 0.98 | 0.90 | | | 0.004 | | | 0.056 | | | 0.89 |

Supplementary Material 6

Table ST3. **Discrimination indices for all ROIs.** Discrimination indices (*D*) and *p-*values for all ROIs, grouped by object, scene, hippocampal, and other ROIs. Shown are the coarse, mid, and fine discriminations for encoding (red), recall (blue), and cross-discrimination (purple). The *p*-threshold indicates the FDR cutoff for significance, and significance is colored based on the key.

| **Scenes /**  **Objects** | | | Encoding | | Recall | | Cross | |  |  |  |  |  |  | | |  | |  |  |  |  |  |
| --- | --- | --- | --- | --- | --- | --- | --- | --- | --- | --- | --- | --- | --- | --- | --- | --- | --- | --- | --- | --- | --- | --- | --- |
|  |  |  | ***D*** | *p* | ***D*** | *p* | ***D*** | *p* | ***p-threshold*** | |  |  | **Key** | | | | | | | | | |  |
| Obj. ROIs | | LO | **0.084** | 2.0E-5 | **0.011** | 0.001 | **0.002** | 0.23 | 0.009 |  |  |  |  |  |  |  |  |  |  |  |  |  |  |
|  |  | pFs | **0.193** | 3.0E-5 | **0.027** | 0.003 | **0.043** | 0.006 | 0.012 |  |  |  | Significant Encoding Discriminability (FDR-corrected) | | | | | | | | | |  |
| Sce. ROIs | | PPA | **0.152** | 3.3E-7 | **0.032** | 3.4E-4 | **0.014** | 0.036 | 0.011 |  |  |  | Significant Recall Discriminability (FDR-corrected) | | | | | | | | | |  |
|  |  | MPA | **0.142** | 2.9E-6 | **0.026** | 0.006 | **0.039** | 1.6E-4 | 0.008 |  |  |  | Significant Cross Discriminability (FDR-corrected) | | | | | | | | | |  |
|  |  | OPA | **0.115** | 1.3E-6 | **0.025** | 0.005 | **-0.003** | 0.67 | 0.005 |  |  |  | *p* < 0.05 (does not pass FDR correction) | | | | | | | | | |  |
| Hipp. | | HB | **0.013** | 3.3E-4 | **-0.002** | 0.82 | **0.003** | 0.082 | 3.3E-4 |  |  |  |  |  |  |  | |  | | |  |  |  |
|  |  | T | **0.016** | 0.108 | **0.000** | 0.48 | **0.008** | 0.10 | None |  |  |  |  |  |  |  | |  | | |  |  |  |
| Other | | PRC | **0.039** | 5.7E-5 | **0.013** | 0.004 | **0.018** | 2.5E-4 | 0.004 |  |  |  |  |  |  |  | |  | | |  |  |  |
|  |  | PHC | **0.074** | 4.6E-6 | **0.014** | 0.003 | **0.024** | 4.0E-4 | 0.003 |  |  |  |  |  |  |  | |  | | |  |  |  |
|  |  | EVC | **0.080** | 4.4E-7 | **0.007** | 0.004 | **-0.010** | 0.99 | 0.008 |  |  |  |  |  |  |  | |  | | |  |  |  |
| **Objects** | | | Big vs. Small | | | | | | Tools vs. Non-tools | | | | | | Object Individuation | | | | | | | | |
|  |  |  | Encoding | | Recall | | Cross | | Encoding | | Recall | | Cross | | Encoding | | | Recall | | | | Cross | |
|  |  |  | ***D*** | *p* | ***D*** | *p* | ***D*** | *P* | ***D*** | *p* | ***D*** | *p* | ***D*** | *p* | ***D*** | *P* | | ***D*** | | | *p* | ***D*** | *p* |
| Obj. ROIs | | LO | **0.006** | 0.15 | **-0.001** | 0.56 | **-0.002** | 0.73 | **0.021** | 0.005 | **0.009** | 0.15 | **0.002** | 0.28 | **0.054** | 1.7E-4 | | **-0.017** | | | 0.70 | **0.006** | 0.20 |
|  |  | pFs | **0.005** | 0.15 | **-0.008** | 0.93 | **-0.004** | 0.91 | **0.014** | 0.012 | **-0.005** | 0.89 | **-2.6E-4** | 0.54 | **0.050** | 1.3E-4 | | **0.008** | | | 0.13 | **0.006** | 0.15 |
| Sce. ROIs | | PPA | **-2.5E-4** | 0.52 | **-0.007** | 0.89 | **-0.001** | 0.61 | **0.006** | 0.16 | **-0.009** | 0.96 | **0.001** | 0.43 | **0.034** | 2.2E-4 | | **0.012** | | | 0.07 | **-0.001** | 0.60 |
|  |  | MPA | **-0.004** | 0.74 | **-0.013** | 0.98 | **-0.001** | 0.60 | **-0.008** | 0.83 | **-0.009** | 0.96 | **-0.005** | 0.79 | **0.020** | 0.053 | | **0.021** | | | 0.09 | **0.012** | 0.048 |
|  |  | OPA | **-0.003** | 0.69 | **-0.007** | 0.89 | **-0.003** | 0.74 | **-0.003** | 0.74 | **-0.012** | 0.95 | **-0.001** | 0.57 | **0.055** | 6.6E-5 | | **0.015** | | | 0.17 | **0.010** | 0.15 |
| Hipp. | | HB | **-0.007** | 0.97 | **-0.007** | 0.95 | **-0.005** | 0.93 | **0.001** | 0.40 | **-0.004** | 0.84 | **0.002** | 0.27 | **0.005** | 0.31 | | **0.018** | | | 0.009 | **-0.002** | 0.61 |
|  |  | T | **-0.008** | 0.91 | **-0.009** | 0.87 | **-0.007** | 0.89 | **-0.004** | 0.70 | **0.002** | 0.38 | **0.000** | 0.53 | **-0.002** | 0.55 | | **0.002** | | | 0.42 | **0.002** | 0.40 |
| Other | | PRC | **-0.005** | 0.78 | **-0.005** | 0.74 | **-0.005** | 0.84 | **0.002** | 0.40 | **-0.004** | 0.75 | **0.002** | 0.24 | **0.010** | 0.18 | | **0.003** | | | 0.36 | **0.004** | 0.35 |
|  |  | PHC | **-0.002** | 0.65 | **-0.003** | 0.76 | **0.001** | 0.38 | **0.001** | 0.47 | **-0.003** | 0.76 | **0.000** | 0.51 | **0.005** | 0.34 | | **0.003** | | | 0.35 | **0.001** | 0.42 |
|  |  | EVC | **0.007** | 0.13 | **-0.006** | 0.87 | **-0.005** | 0.95 | **-0.002** | 0.63 | **0.001** | 0.35 | **0.003** | 0.22 | **0.114** | 5.1E-7 | | **0.023** | | | 0.008 | **-0.022** | 0.99 |
| **Scenes** | | | Open vs. Closed | | | | | | Natural vs. Manmade | | | | | | Scene Individuation | | | | | | | | |
|  |  |  | Encoding | | Recall | | Cross | | Encoding | | Recall | | Cross | | Encoding | | | Recall | | | | Cross | |
|  |  |  | ***D*** | *p* | ***D*** | *p* | ***D*** | *p* | ***D*** | *p* | *D* | *P* | ***D*** | *p* | ***D*** | *p* | | ***D*** | | | *p* | ***D*** | *p* |
| Obj. ROIs | | LO | **0.010** | 0.009 | **-0.006** | 0.82 | **0.004** | 0.25 | **0.031** | 0.001 | **-0.011** | 0.90 | **-0.010** | 0.97 | **0.015** | 0.041 | | **0.012** | | | 0.11 | **-0.007** | 0.69 |
|  |  | pFs | **0.014** | 0.002 | **-0.003** | 0.64 | **0.002** | 0.29 | **0.023** | 0.022 | **-0.003** | 0.63 | **0.002** | 0.38 | **0.020** | 0.035 | | **0.026** | | | 0.009 | **0.003** | 0.37 |
| Sce. ROIs | | PPA | **0.017** | 0.001 | **-0.003** | 0.75 | **0.002** | 0.32 | **0.030** | 0.003 | **0.000** | 0.52 | **0.009** | 0.039 | **0.022** | 7.6E-4 | | **0.023** | | | 0.011 | **0.000** | 0.49 |
|  |  | MPA | **0.010** | 0.029 | **0.002** | 0.39 | **-0.001** | 0.69 | **0.032** | 4.8E-4 | **0.002** | 0.39 | **0.003** | 0.36 | **0.012** | 0.049 | | **0.029** | | | 0.008 | **0.006** | 0.28 |
|  |  | OPA | **0.005** | 0.16 | **-0.003** | 0.65 | **-0.006** | 0.95 | **0.018** | 0.06 | **-0.004** | 0.71 | **0.003** | 0.25 | **0.024** | 0.002 | | **0.024** | | | 0.050 | **0.004** | 0.35 |
| Hipp. | | HB | **0.001** | 0.42 | **0.004** | 0.25 | **-0.003** | 0.76 | **0.001** | 0.44 | **-0.002** | 0.65 | **0.003** | 0.28 | **0.007** | 0.22 | | **-0.001** | | | 0.55 | **-0.010** | 0.93 |
|  |  | T | **0.013** | 0.018 | **0.008** | 0.06 | **0.006** | 0.16 | **0.007** | 0.14 | **-0.004** | 0.66 | **0.001** | 0.42 | **-0.024** | 0.97 | | **0.009** | | | 0.24 | **-0.011** | 0.85 |
| Other | | PRC | **0.010** | 0.031 | **-0.001** | 0.57 | **-3.7E-5** | 0.50 | **0.008** | 0.26 | **-0.006** | 0.81 | **0.002** | 0.41 | **-0.016** | 0.83 | | **0.007** | | | 0.26 | **-0.006** | 0.70 |
|  |  | PHC | **0.012** | 0.014 | **0.003** | 0.25 | **0.003** | 0.23 | **0.010** | 0.11 | **-0.003** | 0.70 | **0.002** | 0.37 | **0.004** | 0.34 | | **0.014** | | | 0.13 | **-0.006** | 0.81 |
|  |  | EVC | **0.004** | 0.21 | **-0.008** | 0.91 | **-1.2E-4** | 0.51 | **0.012** | 0.052 | **-0.012** | 0.98 | **-0.006** | 0.91 | **0.023** | 0.005 | | **0.020** | | | 0.06 | **-0.011** | 0.90 |

Supplementary Material 7: ROI-to-ROI similarity analysis

**Methods**

We investigated ROI-to-ROI correlations to assess similarity between region of interest (ROI) representations separately during encoding and recall. For each participant, we constructed a representational similarity matrix (RSM) for each ROI, as described in the Main Methods section entitled *Representational Similarity Analyses and Discrimination Indices*. We then constructed an ROI-to-ROI similarity matrix by conducting secondary correlations between the RSMs for each pair of ROIs. Specifically, we correlated the ROI RSM calculated with half of the runs with the one created using the other half of the runs, so we could also look at an ROI’s similarity to itself across runs and avoid any common fluctuations within runs (Henriksson et al. 2015). We additionally conducted multidimensional scaling (MDS), to visualize the similarity structure of these ROIs in two dimensions. This ROI-to-ROI similarity analysis was conducted separately for encoding trials and recall trials.

**Results and Discussion**

We looked at correlations of the RSMs between each pair of ROIs, separately for objects and scenes during both encoding and recall (Figure SF3). For encoding objects and scenes, all regions were significantly self-correlated across runs (all *p* < 0.005), indicating that these regions show consistent patterns across runs and exemplars. Within the MDS for encoding both objects and scenes, scene- and object-selective regions are grouped together, with object regions LO and pFs similar to each other for object encoding, and scene regions PPA, MPA, OPA similar to each other for scene encoding. Meanwhile, hippocampal regions are highly dissimilar from all scene- and object-selective regions.

During recall, the representational structure of these regions appears to transform. All ROIs are still significantly correlated with themselves for recalling objects and scenes (all *p* < 0.05). These significant within-region correlations during both encoding and recall suggest there may be similarities in representational structure between encoding and recall that do not necessarily align with the different levels of information we specifically examined. For example, there may be some structure at the level of an individual stimulus (e.g., if one stimulus is dissimilar from all others), but this may not result in discriminability of object category or mid-level information (e.g., whether it is a tool or not). During recall, scene- and object-selective regions are more dissimilar from each other, and these regions and hippocampus are more similar (Figure SF3). To quantify cortical-hippocampal similarity, we compared the mean ranked correlation coefficient of the scene- and object-selective ROIs (pFs, LO, PPA, MPA, OPA) with the hippocampal ROIs (Hip-HB, Hip-T) between encoding and recall. In a 2-way ANOVA for stimulus class (object/scene) and memory process (encoding/recall), there was a significant difference in cortical-hippocampal similarity between encoding and recall (*F*(1,84) = 8.40, *p* = 0.009), but no main effect of stimulus category (*F*(1,84) = 3.30, *p* = 0.084) nor an interaction (*F*(1,84) = 0.37, *p* = 0.549). A post-hoc independent samples t-test indicated that cortical-hippocampal similarity was significantly higher during recall than encoding (*t*(43) = 2.34, *p* = 0.024). To test whether these changes in similarity merely reflected added noise during recall, we looked at whether within-ROI correlations (which can also be considered a measure of signal reliability) changed between encoding and recall. Visual areas showed decreased within-ROI similarity between encoding and recall for both object stimuli (paired t-test, *t*(21) = 3.59, *p* = 0.002) and scenes (*t*(21) = 3.86, *p* = 9.12 × 10^-4^). However, hippocampal areas showed no change in within-ROI similarity between encoding and recall for objects (*t*(21) = 1.32, *p* = 0.202) and scenes (*t*(21) = -0.83, *p* = 0.417). These results suggest that the patterns during recall do not reflect a general effect of noise across the brain.

Thus, across both objects and scenes, representations in scene- and object-selective cortex and hippocampus are more similar during recall than encoding. This could reflect a transformation of the representational structure itself between encoding and recall. Future studies will need to disentangle the nature of these apparent representational transformations.

Supplementary Material 8

**Figure SF3** – **Comparison of stimulus representations between regions.** Matrices showing Pearson correlations between the RSMs of every ROI pair, using separate halves of the data. Matrices for encoding and recall are shown separately for objects and scenes, and are ranked for visualization purposes. In the matrices, dark blue indicates the most similar ROIs, while white indicates the most dissimilar ROIs. ROIs can be conceptually divided into four groups as indicated by the colored squares – early visual areas (red: EVC), object-selective areas (orange: LO, pFs), scene-selective areas (green: PPA, MPA, OPA), and hippocampus (blue: Hip-HB, Hip-T). Next to the encoding and recall RSMs are multidimensional scaling (MDS) plots, which transform a given RSM into a two-dimensional representation where shorter distances indicate higher similarity. With this, one can visualize which ROIs have similar representations to another. During encoding, while hippocampal regions (blue) are very distinct in their representations from visual regions, they become more similar to visual regions during recall.
